# IFN-γ primes bone marrow neutrophils to acquire regulatory functions in severe viral respiratory infections

**DOI:** 10.1101/2023.11.23.568456

**Authors:** Florent Creusat, Youenn Jouan, Loïc Gonzalez, Emilie Barsac, Guy Ilango, Roxane Lemoine, Daphnée Soulard, Chloé Boisseau, Antoine Guillon, Qiaochu Lin, Carolina de Amat Herbozo, Valentin Sencio, Nathalie Winter, François Trottein, Mustapha Si-Tahar, Benoit Briard, Thierry Mallevaey, Christelle Faveeuw, Thomas Baranek, Christophe Paget

## Abstract

Neutrophil subsets endowed with regulatory/suppressive properties are widely regarded as deleterious immune cells that can jeopardize antitumoral response and/or antimicrobial resistance. Here, we describe a sizeable fraction of neutrophils characterized by the expression of Programmed death-ligand 1 (PD-L1) in biological fluids of humans and mice with severe viral respiratory infections (VRI). Biological and transcriptomic approaches indicated that VRI-driven PD-L1^+^ neutrophils are endowed with potent regulatory functions and reduced classical antimicrobial properties, as compared to their PD-L1^-^ counterpart. VRI-induced regulatory PD-L1^+^ neutrophils were generated in the bone marrow in an IFN-γ-dependent manner and were quickly mobilized into the inflamed lungs where they fulfilled their maturation. Neutrophil depletion and PD-L1 blockade during experimental VRI resulted in higher mortality, increased local inflammation and reduced expression of resolving factors. These findings suggest that PD-L1^+^ neutrophils are important players in disease tolerance by mitigating local inflammation during severe VRI and that they may constitute relevant targets for future immune interventions.

## Introduction

Severe VRI is a clinical condition characterized by the rapid onset of diffuse inflammation in lung parenchyma. Among common respiratory viruses in severe pneumonia are influenza A virus (IAV), influenza B virus, rhinoviruses and coronaviruses. Viral pneumonia triggers a complex and multifaceted host response involving discrete immune cell populations as well as many inflammatory soluble factors that exert protective or deleterious functions according to the severity and/or the course of infection^1^. In worse clinical presentations, pneumonia can progress into life-threatening acute respiratory distress syndrome (ARDS) characterized by a sustained and dysregulated immune response^2,3^. Why are some individuals more or less prone to develop ARDS? A current hypothesis is the advantage of a complex immune trait referred to as “resilience”. This could be defined by the ability of the host to mount a sufficient response to allow pathogen containment/elimination (“resistance”) while preventing over-inflammation and preserving tissue integrity (“tolerance”)^4^. During viral ARDS, severe clinical phenotypes are mainly attributable to a defective immune tolerance that can culminate in fatal immunopathology^5^.

In this paradigm, neutrophils have been associated with poor prognosis in severe viral pneumonia/ARDS^6^ and neutrophil dysfunction has recently emerged as a potential cause of death in SARS-CoV-2^7–9^ and experimental IAV infections^10^. These deleterious effects rely on the release of neutrophilic factors such as reactive oxygen species and neutrophil extracellular traps^11–13^, which contribute to sustain the local inflammation and tissue damages. In addition, neutrophils can also participate in the resolution of inflammation, tissue healing/regeneration and help in antibody production^14–16^. Whether such functional diversity occurs during acute viral infection is unknown.

During viral pneumonia, influx of neutrophils in the lung tissue has been shown to rely on an increased neutrophil egress from the bone marrow (BM), a process referred to as “emergency granulopoeisis”^17^, enabling to maintain their presence in inflamed tissues. This “on-demand” mechanism relies on enhanced proliferation of myeloid precursors triggered by soluble factors (*e.g.* cytokines and/or growth factors) and/or pathogen-associated molecular patterns^17^. However, whether these factors can more finely influence neutrophil differentiation/functions remain poorly described during viral pneumonias.

By combining clinical data and experimental IAV-induced severe pneumonia, we identified a subset of regulatory neutrophils characterized by the expression of Programmed death-ligand 1 (PD-L1). During viral pneumonia, the emergence of this subset appeared to be remotely imprinted in the BM in an IFN-γ-dependent manner prior to their migration to the inflammatory site. This leads to a transcriptional program associated with regulatory properties. Importantly, both neutrophil depletion and PD-L1 blockade heightened susceptibility to experimental viral pneumonia. Thus, our study highlights an important IFN-γ/PD-L1^+^ neutrophil axis serving as a feedback loop to control inflammation during severe viral pneumonia. We bring to light an immunological concept in which viral pneumonia remotely controls BM neutrophil differentiation process to assign them with regulatory functions that may pave the way to new therapeutic options in viral pneumonia and ARDS.

## Results

### PD-L1-expressing neutrophil accumulation correlates with disease severity in severe VRI patients

To analyse the phenotype of neutrophils during severe VRI, we enrolled 35 patients admitted in intensive care unit for pneumonia/ARDS with diagnosed SARS-CoV-2 (n = 27) or Flu (n = 8) infections. The baseline characteristics of these patients are presented in **Table 1**. The proportion of CD10^low^ immature neutrophils^18^ was increased in the blood of VRI patients as compared to healthy donors (**Fig. 1a, b**) suggesting an active emergency granulopoiesis. In addition, neutrophils from VRI patients expressed elevated levels of the inhibitory costimulatory molecule PD-L1 compared to healthy donors (**Fig. 1a, c**). Notably, PD-L1 expression could be found on both mature (CD10^high^) and immature (CD10^low^) neutrophil subsets (**Fig. 1a**) with similar proportions (**Fig. S1a**). The concentration of circulating PD-L1^+^ neutrophils was slightly higher in VRI patients with ARDS as compared to non-ARDS patients (**Fig. 1d**). The relative proportion of the PD-L1^+^ neutrophils was further increased in endotracheal aspirates (ETA) of intubated patients (n = 23) compared to the blood (**Fig. 1e**) suggesting their accumulation in the airways. Hypoxemia positively correlated with the proportion of airway but not circulating PD-L1^+^ neutrophils (**Fig. 1f and S1b**). Altogether, these data suggest that severe VRI is associated with a local and systemic accumulation of PD-L1-expressing neutrophils.

**Figure 1:**
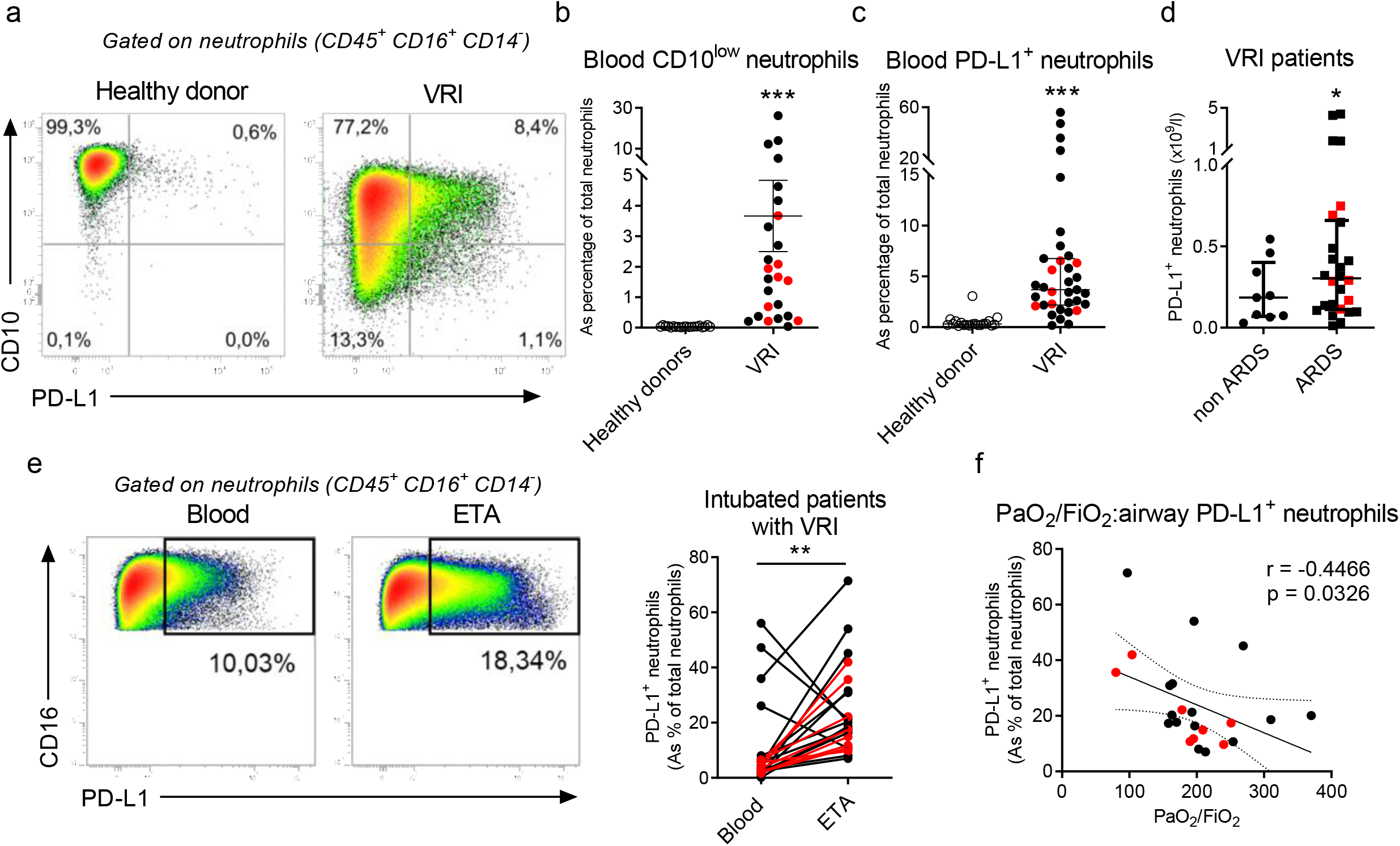
Neutrophils in patients with severe virus-induced ARDS. **a-d**, Flow cytometry analysis of circulating neutrophils in blood of 35 patients with severe viral pneumonia (black dots: COVID-19, red dots: Flu) within 48 hrs post-admission. **a**, Phenotype of circulating neutrophils (CD45^+^ CD3^-^ CD16^+^ CD14^-^). Representative dot plots of CD10 and PD-L1 expression on circulating neutrophils from healthy donors and VRI patients are shown. **b-c**, Relative proportion of immature CD10^low^ (**b**) or PD-L1^+^ (**c**) neutrophils within the total neutrophil compartment in blood of controls and patients. Individual and means ± SEM are shown (n = 20-25/group). **d**, Concentration of circulating PD-L1^+^ neutrophils according to the ARDS status. Individual values and means ± SEM are depicted (n = 9-26/group). **e**, Representative dot plots of PD-L1 expression on neutrophils from blood or endotracheal aspirates (ETA) are shown in the left panel. Paired-analysis of PD-L1^+^ neutrophils in blood and ETA of matched VRI patients. **f**, Spearman’s rank correlation of airway PD-L1^+^ neutrophils and hypoxemia levels on admission of intubated patients with viral pneumonia. ns, not significant; *, p < 0.05; **, p < 0.01; ***, p < 0.001.

### PD-L1^+^ neutrophils accumulate in the lungs during experimental viral pneumonia

To evaluate the biology of lung PD-L1^+^ neutrophils in severe VRI, we infected mice with a highly pathogenic strain of IAV (H3N2 A/Scotland/20/74). In this model, mice die from a deleterious inflammatory response in the lungs despite complete viral clearance^19^, which resembles clinical ARDS. We first assessed the dynamic of airway and parenchyma lung neutrophils during the course of IAV infection. The relative proportion of airway neutrophils rapidly increased upon IAV infection, while it remained unchanged in the lung parenchyma (**Fig. 2a, upper and middle panels**). However, the absolute number of neutrophils was increased in both compartments to peak at 8 days post-infection (dpi) (**Fig. 2a, lower panels**). Similar to VRI patients, neutrophils from IAV-infected mice could be segregated based on PD-L1 expression in both airways and parenchyma (**Fig. 2b, upper panels**). The proportion and absolute number of PD-L1^+^ neutrophils increased as early as 4 dpi (**Fig. 2b, middle and lower panels**) to become the major neutrophil subset during the inflammatory phase (6-10 dpi). Notably, neutrophils accounted for ∼ 50% of total PD-L1-expressing leukocytes in airways at 8 dpi (**Fig. S2**) while constituting a minor fraction in the parenchyma (**Fig. S2**). Together, these data demonstrate that PD-L1^+^ neutrophils are a hallmark of the acute phase of experimental IAV infection in the lung.

**Figure 2:**
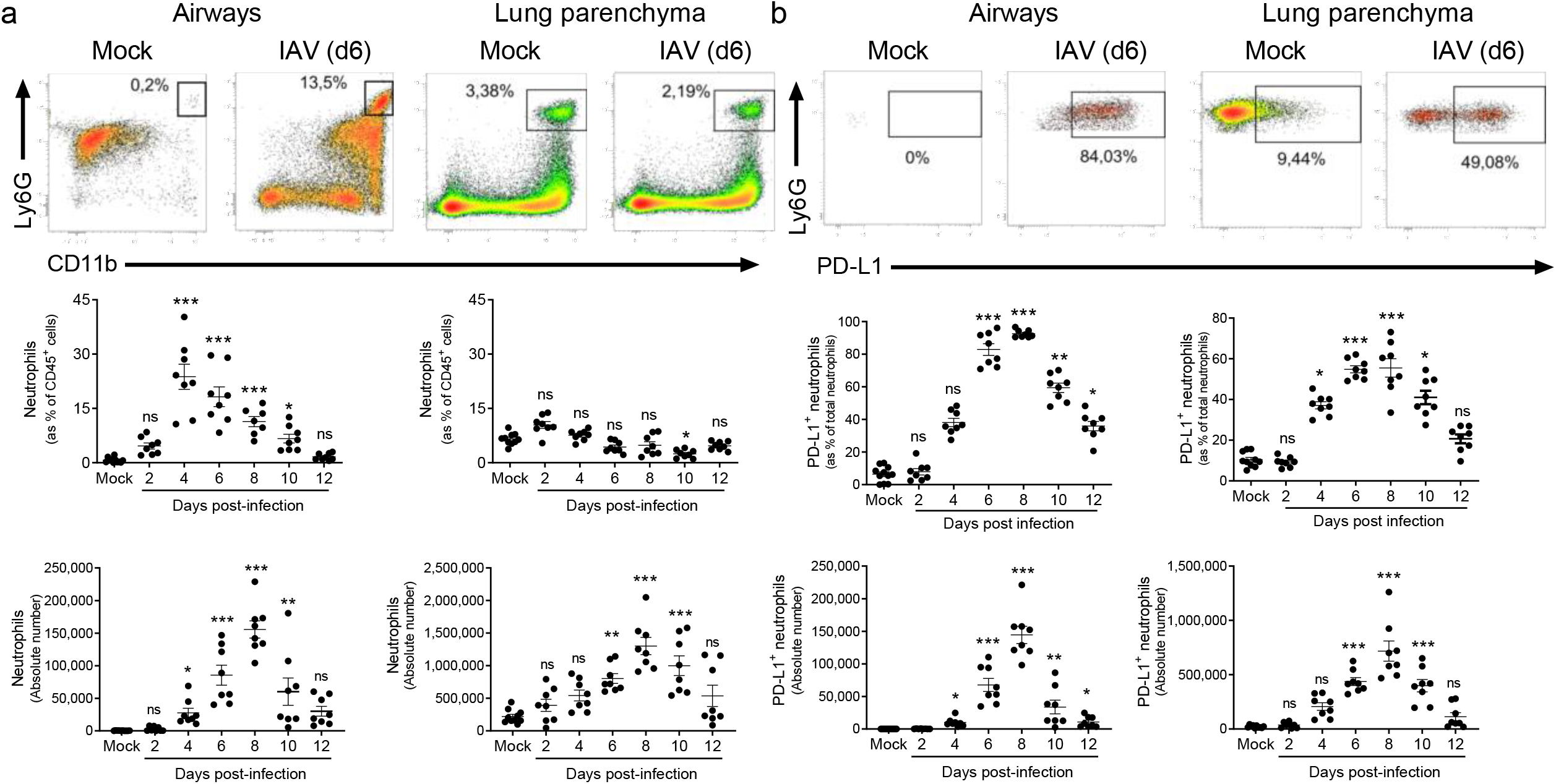
Dynamics and phenotype of neutrophils during experimental IAV infection. **a-b**, WT C57BL/6j mice were i.n. infected with mock or IAV (150 pfu) A/H3N2/Scotland/20/74 strain. Mice were euthanized at indicated time points and BAL and lungs were harvested. **a**, Relative proportion and absolute numbers of neutrophils in airways and lung parenchyma were evaluated by flow cytometry. Representative dot plots of neutrophils in airway and lung parenchyma from mock and IAV-infected mice (6 dpi) are shown in the upper panel. Individual and means ± SEM pooled from three independent experiments are shown in the lower panel (8-10 mice/group). **b**, Relative proportion and absolute numbers of PD-L1^+^ neutrophils in airways and lung parenchyma were evaluated by flow cytometry. Representative dot plots of PD-L1^+^ neutrophils in airway and lung parenchyma from mock and IAV-infected mice (6 dpi) are shown in the upper panel. Individual and means ± SEM pooled from three independent experiments are shown in the lower panel (8-10 mice/group). ns, not significant; *, p < 0.05; **, p < 0.01; ***, p < 0.001.

### Lung PD-L1^+^ and PD-L1^-^ neutrophils display discrete transcriptional signatures

To characterize these two neutrophil subsets, we profiled the transcriptomes of PD-L1^+^ and PD-L1^-^ neutrophils isolated from the lungs of IAV-infected mice using single-cell (sc) capture using droplet-based system (**Fig. 3a**). Following established quality controls and filtering steps^20^, 2,999 cells (1,834 PD-L1^+^; 1,165 PDL-1^-^) were used for downstream analyses. Cell clustering was defined using the Louvain algorithm^21^ after merging datasets and reducing dimensionality of the merged dataset through Uniform Manifold Approximation and Projection (UMAP). Analysis of neutrophil marker genes (*S100a9, Il1b, Dusp1, Lgals3*) confirmed the lineage specificity of our datasets **(Fig. S3a**). We identified five neutrophil clusters (**Fig. 3b**) with variable prevalence (**Fig. S3b**). PD-L1^-^ neutrophils belonged almost exclusively to clusters 1 and 3 (**Fig. 3c and Fig. S3c**), whereas PD-L1^+^ neutrophils were distributed among clusters 0, 2 and 4 (**Fig. 3c and Fig. S3c**). This suggested substantial transcriptional differences between the two subsets as well as intra-subset heterogeneity (**Fig. 3c and Table S1**).

**Figure 3:**
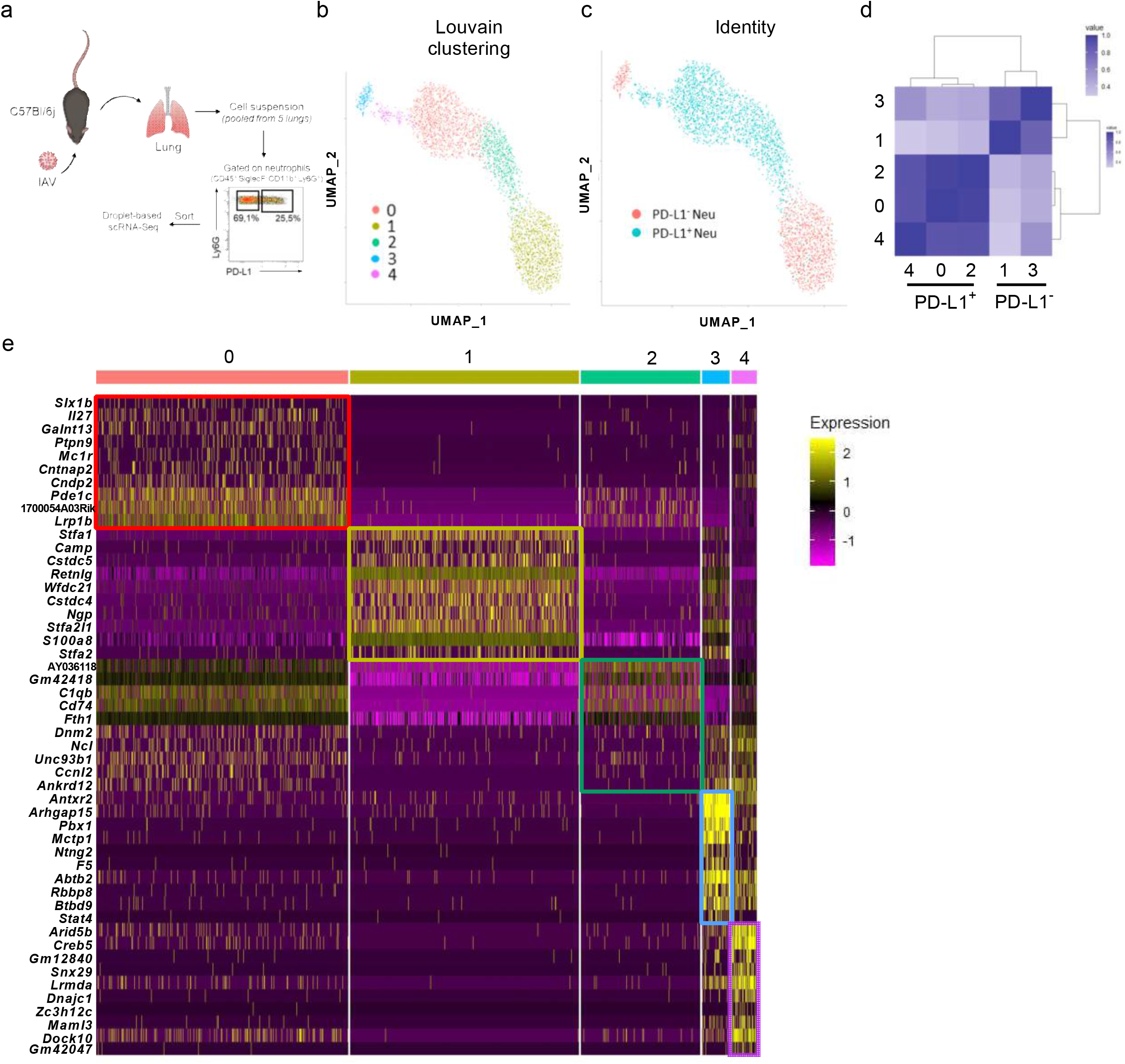
scRNA-seq profiling of lung neutrophil subsets from IAV-infected mice. **(a-e),** WT C57BL/6j mice were i.n. instilled with IAV (150 pfu) A/H3N2/Scotland/20/74 strain and euthanized at day 11. Lungs were collected and processed prior neutrophil sorting based on PD-L1 expression. **a**, Representation of the experimental workflow used to generate scRNA-seq data. **b**, Identification of cell clusters using the graph-based Louvain algorithm (resolution = 0.5) on Uniform Manifold Approximation and Projections (UMAP). Each dot represents one cell (2,999 cells). **c**, Identity of PD-L1^+^ and PD-L1^-^ neutrophils projected on the UMAP. **d**, Clustermap of neutrophil subsets comparing each pair of clusters using the Pearson’s correlation with hierarchical clustering. **e**, Expression of the top-10 marker genes for each cluster identified in (**b**).

To assess further the transcriptional relationships between subsets/clusters, we generated a clustermap using the Pearson’s method to order clusters by linkages based on similarity between the correlations (**Fig. 3d**). Clusters 0, 2 and 4 (PD-L1^+^ neu) were closely related with highest similarities between clusters 0 and 2 (**Fig. 3d**). Regarding PD-L1^-^ clusters, clusters 1 and 3 appeared to be related albeit with lower correlation than within PD-L1^+^ neutrophil-associated clusters (**Fig. 3d**).

Comparative analysis of gene expression between the five clusters (**Fig. 3e and Table S1**) indicated that clusters 0, 2 and 4 (PD-L1^+^ neu) shared many differentially expressed genes (DEGs) including genes encoding for MHC-II (and -associated) molecules such as *H2-Aa*, *Cd74*, *H2-Ab1*, *H2-Eb1, H2-K1* and *H2-D1* (**Fig. 3e and Table S1**). Among other notable DEGs, PD-L1^+^ clusters were also defined by high expression of many transcripts encoding for chemokines that have been shown to be released by suppressive tumor-associated neutrophils^22^ such as *Ccl2*, *Ccl3*, *Ccl4*, *Ccl5, Ccl12*, *Ccl17* and *Cxcl16* (**Table S1**). Thus, the transcriptomes of PD-L1^+^ neutrophils suggest a specialised subset with regulatory/suppressive functions. PD-L1^-^ neutrophils (clusters 1 and 3) presented several DEGs encoding for classical markers of neutrophils as *S100a8, S100a9, Il1b, Cxcl2, Ccl6* and *Jund* (**Fig. 3e and Table S1**). Of note, numerous DEGs (*Chil3, Retnlg, Ngp, Camp*) in clusters 1 and 3 (**Fig. 3e and Table S1**) were reminiscent of differentiating BM neutrophils^20^. Moreover, PD-L1^-^ neutrophils preferentially expressed a gene signature of neutrophils with classical antimicrobial (**Fig. S3d**) and phagocytic (**Fig.S3e**) activities.

Clusters 3 (PD-L1^-^) and 4 (PD-L1^+^) were characterized by a high median number of genes, compared to all other clusters (**Fig. S3f**), reminiscent of a strong transcriptional activity associated with cell differentiation and immaturity. In line, these two clusters expressed specific signatures associated with cytoskeleton, gene expression and cell cycle (**Fig. S3g**).

This analysis reveals that PD-L1^-^ and PD-L1^+^ neutrophils have distinct transcriptional signatures during IAV infection, which are evocative of classical inflammatory and regulatory/suppressive functions, respectively.

### PD-L1^+^ neutrophils are immature and apoptosis-resistant

To evaluate their level of maturation, we first performed microscopy-based morphological analysis on isolated lung neutrophil subsets from IAV-infected mice. Interestingly, the PD-L1^+^ fraction was enriched for metamyelocytes and banded neutrophils, whereas the PD-L1^-^ fraction contained a vast majority of classical mature neutrophils with uniform morphology (**Fig. 4a**). Higher prevalence of immature stages within the PD-L1^+^ fraction could also be illustrated based on CD101 expression^23^ (**Fig. 4b**). In addition, a large proportion of PD-L1^+^ neutrophils from IAV-infected mice were FSC^high^ (**Fig. S4a**), which is reminiscent of an immature phenotype^24^. The presence of immature stages in the PD-L1^+^ subset could also be illustrated by the higher proportion of Ki67^+^ cells (**Fig. 4c**) as well as a higher transcriptional G2M cell cycle score in PD-L1^+^ neutrophil-associated clusters (**Fig. S4b**). To finely map the maturation stages at the transcriptional level, we adapted to our datasets the “neutrotime” signature^20^, a model that enables to project neutrophils onto a single maturation ordering from BM precursors to mature neutrophils in peripheral tissues under normal and inflamed conditions. We took advantage of a published gene set^23^ to identify neutrophil precursors within our dataset. A module score for each cell was calculated and revealed a high expression of the precursor signature in cluster 3 (**Fig. S4c**). Thus, we calculated a pseudotime^25^ using cluster 3 as a root (**Fig. S4d**). The linear ordering of neutrophils according to the maturation score indicated that cluster 4 comprised immature cells (**Fig. 4d**). Regarding other PD-L1^+^ clusters, the model suggested that cluster 0 was less mature than cluster 2 (**Fig. 4d**). The cluster 1 that encompasses the vast majority of PD-L1^-^ neutrophils emerged as the most mature subset within our dataset (**Fig. 4d**).

**Figure 4:**
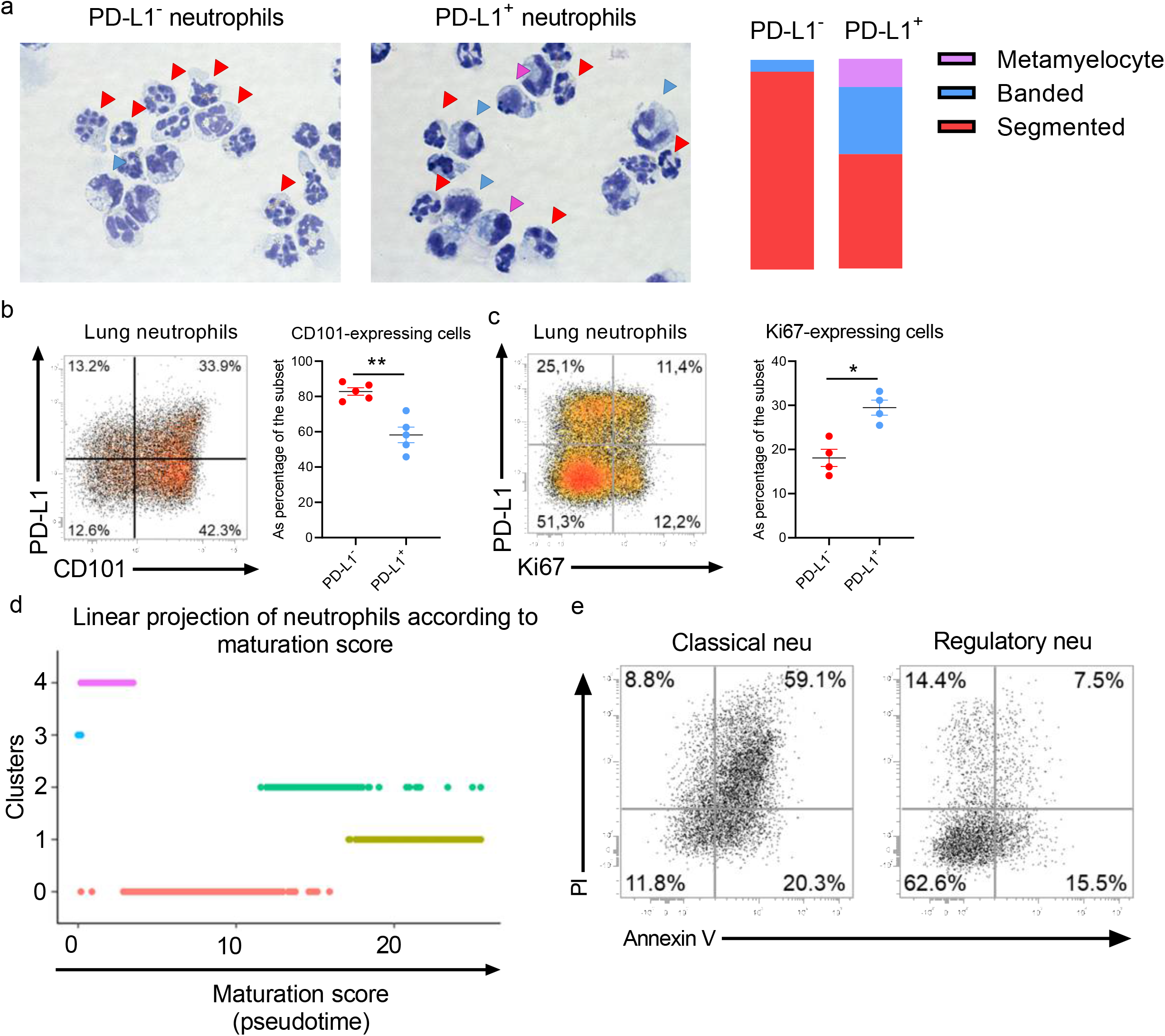
Morphology, maturation and lifespan of lung neutrophilic subsets in IAV-infected mice. **a-e,** WT C57BL/6j mice were i.n. instilled with mock or IAV (150 pfu) A/H3N2/Scotland/20/74 strain. Mice were euthanized and the whole lungs were collected at 7 dpi. **a**, Representative light microscopy pictures of cytospins of purified lung neutrophil subsets. Neutrophil developmental stages were defined based on morphology and depicted in the right panel (n = 3); x 200, original magnification. **b-c**, Representative dot plots of CD101 (**b**) or Ki67 (**c**) expression on lung neutrophils from IAV-infected mice based on PD-L1 expression. Relative proportions of CD101-(**b**) or Ki67-expressing (**c**) cells per subset are shown in the right panel. Individual values and means ± SEM from two independent experiments (4-5 mice/group) are shown. **d**, linear projection of cells according to pseudotime score from Monocle3 according to cluster identities. **e,** Apoptotic profile of lung neutrophil subsets from IAV-infected mice. A representative dot plot of two independent experiments for both subsets is shown (6 mice/group).

In parallel, we also interrogated the potential differential apoptotic profile of the two neutrophil subsets in IAV-infected mice. Based on propidium iodide/Annexin V staining, classical neutrophils expressed a marked phenotype of late-apoptotic cells compared to the regulatory subset (**Fig. 4e**). In line, the neutrophil transcriptomes confirmed an enrichment for genes associated with negative execution of apoptosis in PD-L1^+^ clusters (**Fig. S4e**). However, this signature was also enriched in cells of cluster 3 in agreement with their immature profile (**Fig. S4e**). Moreover, PD-L1^+^ neutrophil-associated clusters displayed increased expression of *Cd47* (**Fig. S4f**), a “don’t eat me” signal that may confer resistance to efferocytosis^26^. Altogether, lung PD-L1^+^ neutrophils from IAV-infected mice displayed multiple features of immature cells and higher resistance to apoptosis.

### IAV-induced lung PD-L1^+^ neutrophils have regulatory functions

We evaluated the phenotype and functions of lung PD-L1^+^ neutrophils from IAV-infected mice. In line with the expression of numerous genes encoding for MHC-II molecules (**Fig. 3e and Table S1**), we confirmed by flow cytometry that lung PD-L1^+^ neutrophils from IAV-infected mice expressed high levels of MHC-II as compared to their PD-L1^-^ counterparts (**Fig. 5a**). We also tested two gene sets from gene ontology associated with the regulation of the inflammatory response on the transcriptomes. We observed a high signature for “positive regulation of the inflammatory response” in mature PDL-1^-^ neutrophils (cluster 1) while featured genes for “negative regulation of the inflammatory response” were enriched in PD-L1^+^ neutrophil-associated clusters especially the cluster 2 (**Fig. 5b**).

**Figure 5:**
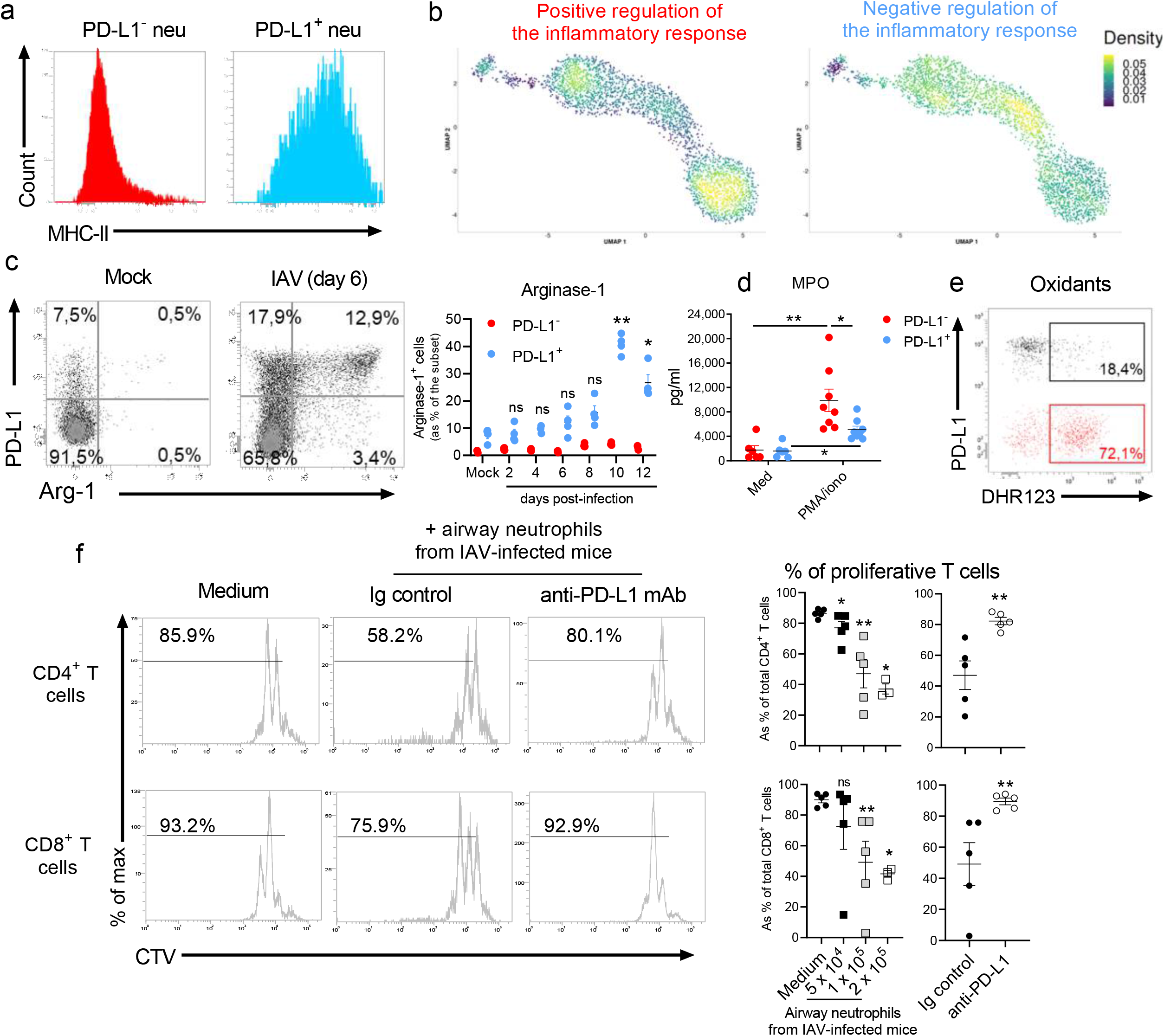
Phenotype and regulatory functions of lung PD-L1^+^ neutrophils. WT C57BL/6j mice were i.n. instilled with mock or IAV (150 pfu) A/H3N2/Scotland/20/74 strain. Mice were euthanized and the whole lungs were collected at indicated time points. **a**, Expression of MHC-II on lung neutrophils from IAV-infected mice (7 dpi) was evaluated by flow cytometry. Representative dot plots according to neutrophilic subset is shown. **b**, Density plot of “positive” and “negative” regulation of the inflammatory response. **c**, Relative proportion of arginase-1-producing neutrophils according to subset and time course of infection. Representative dot plots are shown in the left panel. Individual values and means ± SEM from two independent experiments (4 mice/group/time point) are shown in the right panel. **d**, Levels of MPO produced *ex vivo* by purified neutrophil subsets upon short-term PMA stimulation. Individual values and means ± SEM from two independent experiments are shown. **e**, Levels of oxidants in neutrophil subsets measured 5 min after incubation with DHR123. Representative dot plots from one experiment out of two are shown. *, p < 0.05; **, p < 0.01; ***, p < 0.001. **f**, PD-L1-mediated suppressive effect of neutrophils from IAV-infected mice on T cell proliferation. CTV-stained spleen cells from naive mice were cultured with plate-bound anti-CD3 mAbs with or without indicated numbers of neutrophils from IAV-infected mice (8 dpi) in presence of either Ig control or anti-PD-L1 mAb. Proliferation of CD4^+^ and CD8^+^ T cells was monitored after 72 h based on CTV dilution. Representative dot plots showing CTV expression in CD4^+^ (upper panel) and CD8^+^ (lower panel) T cells are shown in the left panel. Individual values and means ± SEM of proliferation rate from two independent experiments are depicted in the right panel. ns, not significant; *, p < 0.05; **, p < 0.01.

To directly assess the functions of PD-L1^-^ and PD-L1^+^ neutrophils, we first analyzed their production of arginase-1 (Arg-1), a key effector in the regulation of inflammation, which suppresses T cell function^27^. *Arg1* transcripts could be detected in a substantial proportion of cells belonging to PD-L1^+^ neutrophil clusters (**Fig. S5**). Moreover, lung PD-L1^+^ neutrophils produced high amounts of Arg-1 during the course of infection, while PD-L1^-^ neutrophils failed to do so (**Fig. 5c**). Of note, PD-L1^+^ neutrophils displayed a higher capacity to produce Arg-1 during the resolution phase (**Fig. 5c**). Conversely, PD-L1^+^ neutrophils produced less myeloperoxidase (**Fig. 5d**) and oxidants (**Fig. 5e**) than PD-L1^-^ neutrophils.

The potential suppressive activities of PD-L1^+^ neutrophils was then evaluated on activated T cells. Addition of purified neutrophils from airways of IAV-infected mice (containing 90% of PD-L1^+^ neutrophils (**Fig. 2b**)) reduced the proliferative capacities of both CD4^+^ and CD8^+^ T cells in a ratio-dependent manner (**Fig. 5f**). Interestingly, T cell proliferation could be rescued upon PD-L1 blockade (**Fig. 5f**). Together, these data demonstrate that PD-L1^+^ neutrophils that emerge during IAV infection have regulatory/suppressive functions.

### PD-L1^+^ neutrophils are readily detectable in the BM during IAV infection and preferentially migrate towards the lungs

Our data suggested that IAV infection triggers the egress of BM neutrophils containing immature cells (**Fig. 3e and 4b**). We found that the proportion of BM neutrophils rapidly decreased starting at day 4 post-IAV infection (**Fig. 6a, left panel**). Interestingly, although only a minute fraction of PD-L1^+^ neutrophils could be detected in the BM of naive mice, this subset increased as soon as 4 dpi (**Fig. 6a, right panel**). Importantly, BM PD-L1^+^ neutrophils from IAV-infected mice also readily expressed MHC-II (**Fig. S6a**). Consistent with a rapid mobilization of BM neutrophils, the relative proportion of neutrophils increased in the blood of IAV-infected mice (**Fig. 6b, left panel**). Moreover, this was also accompanied with an increase in the proportion of circulating PD-L1^+^ neutrophils (**Fig. 6b, right panel**). In addition, BM and blood PD-L1^+^ neutrophils preferentially expressed CD49d (**Fig. 6c**), a key integrin in the homing of neutrophils in the lung tissue upon infection^28^. Importantly, higher levels of CD49d were also detected on lung PD-L1^+^ neutrophils as compared to their PD-L1^-^ counterpart (**Fig. S6b**). In addition to its anti-apoptotic signals, *Cd47*, which expression is increased in lung PD-L1^+^ neutrophil-associated clusters (**Fig. S4f**), supports the migration of neutrophils to the site of infection^29^. Consistent with a preferential migration of BM PD-L1^+^ neutrophils towards the lung, no enrichment was observed in other peripheral organs of IAV-infected mice such as spleen and liver (**Fig. S6c**), although we noticed a high proportion of PD-L1^+^ neutrophils in the liver of naive mice (**Fig. S6d**).

**Figure 6:**
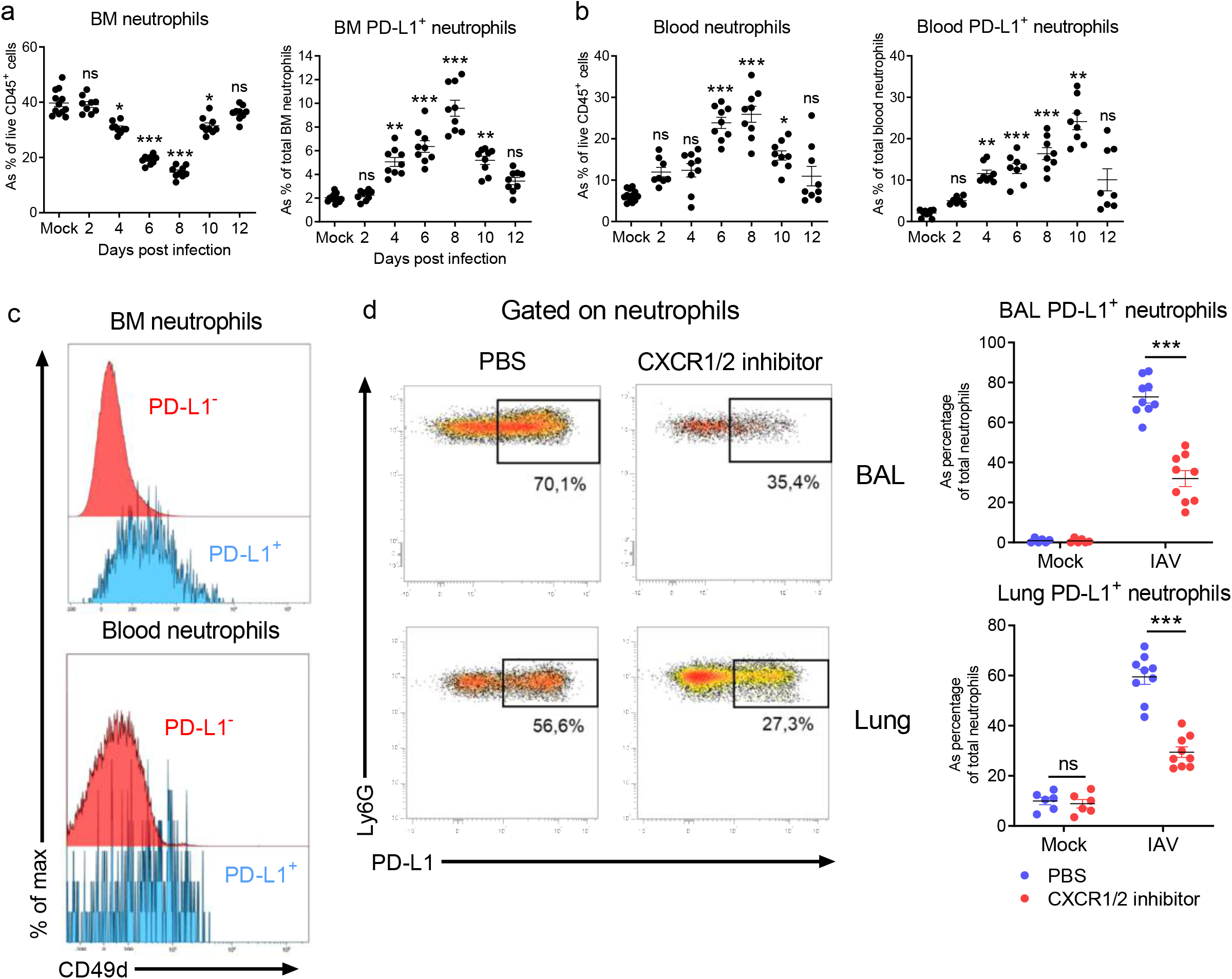
BM origin of IAV-induced PD-L1^+^ neutrophils. **(a-d)**, WT C57BL/6J mice were i.n. infected with IAV (150 pfu) A/H3N2/Scotland/20/74 strain. Mice were euthanized at indicated time points. **a**, Relative proportion of neutrophils (subsets) in BM of IAV-infected mice were evaluated by flow cytometry. Individuals and means ± SEM from three independent experiments are shown (8-12 mice/group). **b**, Frequency of circulating neutrophils (subsets) of IAV-infected mice were evaluated by flow cytometry. Individuals and means ± SEM from three independent experiments are shown (7-12 mice/group). **c**, Expression of CD49d on neutrophil subsets from IAV-infected mice according to PD-L1 expression. Representative histograms of BM (upper panel) and blood (lower panel) neutrophilic subsets are shown. **d**, Effect of CXCR1/2 inhibition on lung PD-L1^+^ neutrophils in IAV-infected mice. Representative dot plots of PD-L1^+^ cells within the BAL (upper panel) and lung parenchyma (lower panel) neutrophil pool from IAV-infected mice treated or not with CXCR1/2 inhibitor are represented in the left panel. Individuals and means ± SEM from two independent experiments are shown (6-9 mice/group) in the right panel. ns, not significant; *, p < 0.05; **, p < 0.01; ***, p < 0.001.

Neutrophil migration from the lung is known to be dependent on CXCR2^30^. Upon CXCR2 blockade^31^, the frequency of lung neutrophils were decreased at 6 dpi (**Fig. S6e**). Importantly, this effect was even more pronounced on the PD-L1^+^ subset (**Fig. 6d**). As expected, this treatment led to an accumulation of neutrophils in the BM (**Fig. S6e**), which was more marked on PD-L1-expressing neutrophils (**Fig. S6f**). Overall, BM PD-L1^+^ neutrophils are quickly mobilized into the lungs during IAV infection.

### IAV-induced IFN-γ programs BM neutrophils for regulatory functions

The emergence of PD-L1^+^ neutrophils in the BM in the early stages of IAV infection suggests that they acquire their regulatory functions during their differentiation and maturation within the BM. First, we interrogated whether the IAV infection influenced the ability of BM haematopoietic stem cells (HSC) to generate neutrophils. Thus, BM HSC from either naive or IAV-infected mice were differentiated *in vitro* into BM-derived neutrophils (BMN). In this setting, both precursors generated the same amount of BMN (**Fig. 7a**) with ∼20% of these latter expressing PD-L1 (**Fig. 7a**).

**Figure 7:**
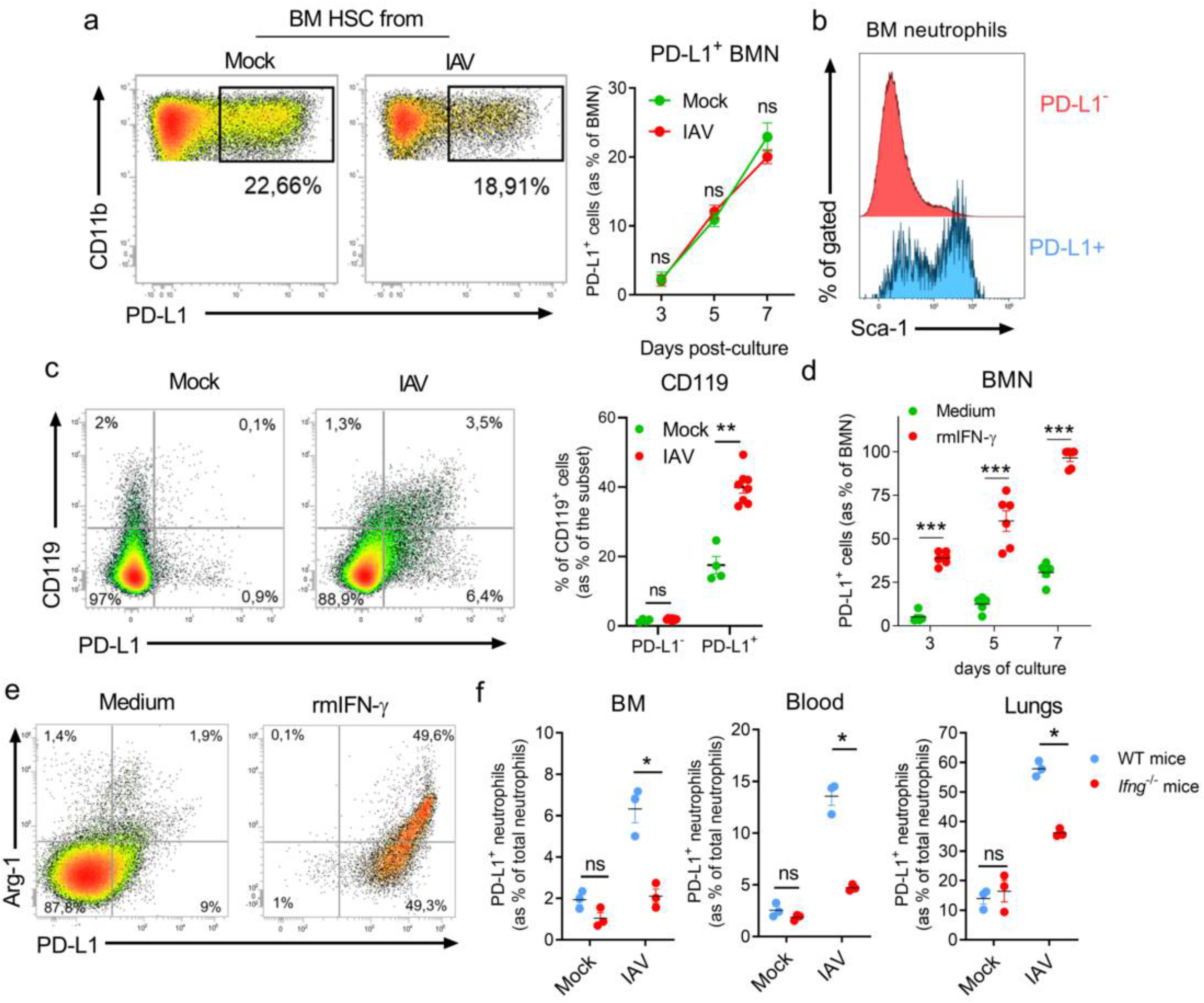
IFN-γ primes BM neutrophils for regulatory functions. **a**, Representative dot plots of the relative proportion of PD-L1^+^ BMN (within total BMN: CD11b^+^ Ly6G^+^) differentiated from stem cells isolated from BM of mock-treated or IAV-infected mice are shown. Means ± SEM at indicated time points from two independent experiments are shown in the right panel. **b**, Representative histograms of Sca-1 expression on BM neutrophil subsets from IAV-infected mice (7dpi). **c**, Expression of CD119 on BM neutrophils according to PD-L1 expression in naive and IAV-infected mice (7 dpi) was evaluated by flow cytometry. Representative dot plots are shown in the left panel. Individuals and means ± SEM of CD119 expression according to neutrophil subset from two independent experiments are shown (4-8 mice/group) in the right panel. **d**, Relative proportions of PD-L1^+^ BMN (within total BMN) differentiated from BM stem cells isolated from naive mice during the course of differentiation in presence of medium, or recombinant mIFN-γ. Individuals and means ± SEM at indicated time points from three independent experiments are shown. **e**, Expression of arginase-1 by BMN subsets differentiated in presence or not of rmIFN-γ. Representative dot plots from two independent experiments are shown. **f**, Relative proportion of PD-L1^+^ neutrophils in various tissues of mock or IAV-infected mice (7 dpi) from WT and *Ifng*^-/-^ mice. Individual values and means ± SEM are shown (3 mice/group). ns, not significant; *, p < 0.05; **, p < 0.01; ***, p<0.001.

Interestingly, BM PD-L1^+^ neutrophils from IAV-infected mice preferentially co-expressed Sca-1 (Ly6A/E) (**Fig. 7b**), a common interferon-stimulated gene (ISG)^32^. In parallel, we detected an increased number of transcripts encoding for IFN-γ in the BM samples of IAV-infected mice as soon as 2 dpi (**Fig. S7a**), which precedes the emergence of PD-L1^+^ neutrophils. Moreover, BM PD-L1^+^ neutrophils from IAV-infected mice preferentially co-expressed IFN-γR1 (CD119) (**Fig. 7c**). Thus, we differentiated BM precursors from naive mice into BMN in the presence of recombinant mouse (rm)IFN-γ. Addition of IFN-γ culminated in the generation of more than 90% of PD-L1^+^ BMN (**Fig. 7d**) co-expressing Sca-1 and MHC-II (**Fig. S7b**), an effect that was abrogated using *Ifngr*^-/-^ BM precursors (**Fig. S7c**). Moreover, PD-L1^+^ BMN produced arginase-1 but only in presence of rmIFN-γ (**Fig. 7e**). Acquisition of the regulatory phenotype was optimal under sustained pressure of IFN-γ (**Fig. S7d**). Notably, the use of rmIFN-γ significantly decreased the absolute number of generated BMN as compared to controls (**Fig. S7e**). This was not associated to a higher mortality (**Fig. S7f**) but to a lower proliferative rate upon rmIFN-γ pressure (**Fig. S7g**) as compared to differentiating PD-L1^-^ BMN. However, this reflected a higher capacity of the PD-L1^-^ subset to cycle as compared to PD-L1^+^ BMN independently from rmIFN-γ usage (**Fig. S7h**) at least under *in vitro* setting. Of note, transcriptomic data also pointed to an early effect of IFN-γ on developing neutrophils since an IFN-γ-mediated signalling pathway signature was enriched in immature lung neutrophil clusters (**Fig. S7i**). Consistent with a role for IFN-γ in the generation of PD-L1^+^ regulatory neutrophils, this signature was also enriched in clusters 0 and 4 as compared to cluster 1 (**Fig. S7i**).

To evaluate the role of IFN-γ *in vivo*, BM PD-L1^+^ neutrophils were analysed in IAV-infected *Ifng*^-/-^ mice. As compared to control mice, a decrease in the number of neutrophils was noted in lungs of *Ifng*^-/-^ mice (**Fig. S7j**). Conversely, neutrophil numbers were increased in the blood of *Ifng*^-/-^ mice (**Fig. S7j**). Analysis of neutrophil subsets indicated that the PD-L1^+^ fraction was the most affected subset upon IFN-γ deficiency in all compartments (**Fig. 7f**). Moreover, remaining PD-L1^+^ neutrophils from *Ifng*^-/-^ mice co-expressed lower levels of Sca-1 and MHC-II as compared to their WT counterparts (**Fig. S7k**). Overall, IAV infection-dependent IFN-γ primes developing neutrophils in the BM resulting in emergence of PD-L1^+^ neutrophils with a regulatory phenotype.

### Presence of PD-L1^+^ neutrophils is associated with protection during IAV infection

Neutrophils have been largely described as deleterious cells in experimental models of IAV infection^10,13,33^. In agreement with this, maintaining neutrophil depletion by means of NIMP-R14 during the entire course of infection led to an almost complete protection associated with limited weight loss and mortality (**Fig. 8a**). Conversely, starting neutrophil depletion on 4 dpi, which corresponds to PD-L1^+^ emergence in the lungs, delayed recovery (prolonged weight loss) and increased mortality (**Fig. 8b**). Importantly, this treatment was not associated with a reduced viral clearance as compared to controls (**Fig. S8a**). However, at 11 dpi, we detected increased levels of soluble inflammatory mediators (IFN-γ, IL-1β and TNF-α) (**Fig. 8c**). CCR2-expressing myeloid cells including monocyte-derived dendritic cells (MoDC) and inflammatory monocytes have been associated to immunopathology during IAV infection^34–36^. Interestingly, a higher infiltration of MoDC and inflammatory monocytes in the lung tissue could be observed in neutrophil-depleted mice (**Fig. 8d and Fig. S8b**). This was accompanied by a reduced number of alveolar macrophages (AM) (**Fig. 8d**), which contribute to tissue repair during IAV infection^37,38^. No change was observed for total conventional DC (**Fig. 8d and Fig. S8b**).

**Figure 8:**
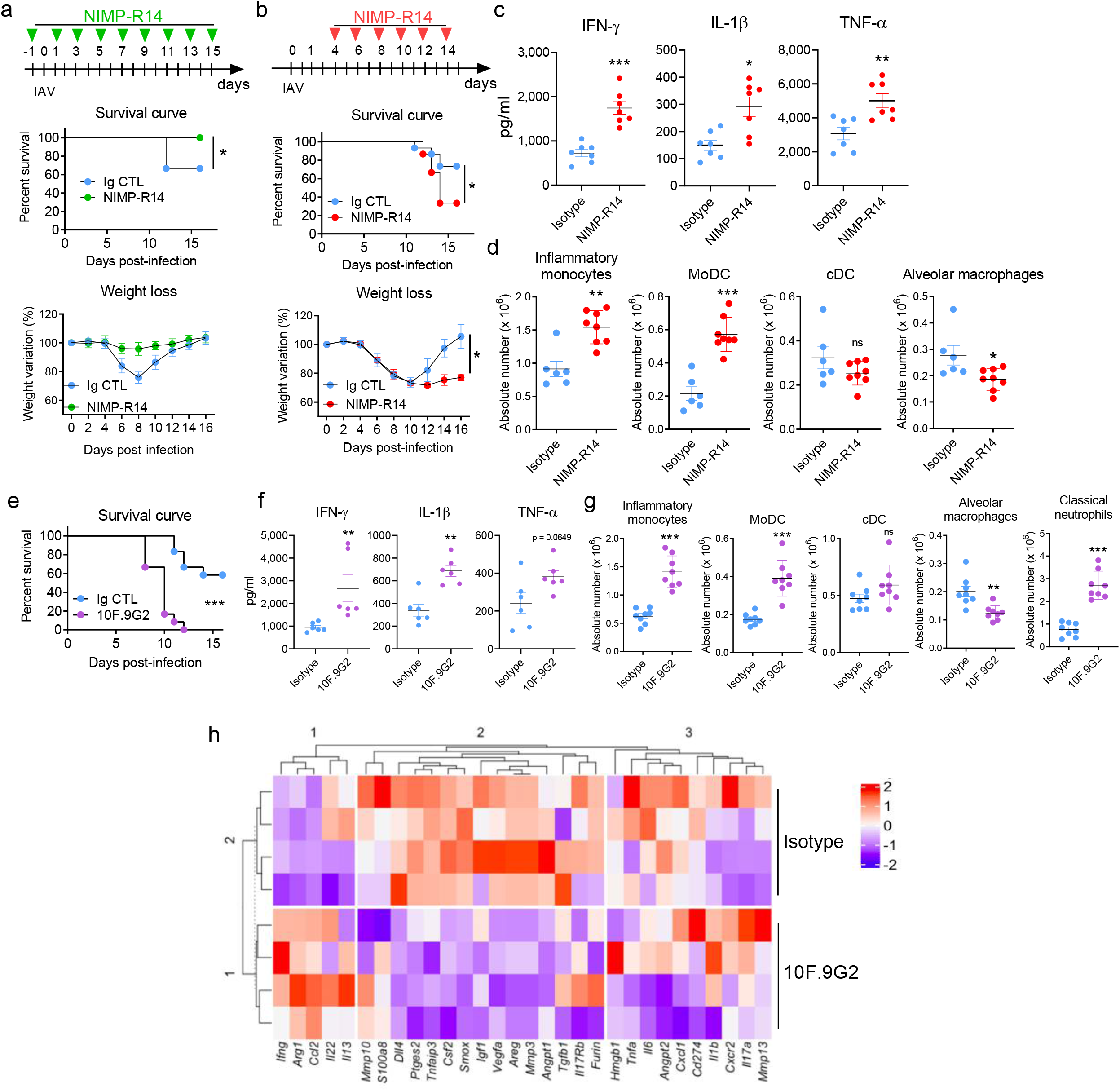
Late neutrophil depletion or PD-L1 blockade influence mouse survival and inflammatory response during IAV infection. **(a-h),** WT C57BL/6j mice were i.n. infected with IAV (150 pfu) A/H3N2/Scotland/20/74 strain. **a**, Mice were treated from -1 dpi and every second day with isotype control or NIMP-R14. Survival (upper panel) and weight loss (lower panel) was monitored daily (12 mice/group). **b**, Mice were treated from 4 dpi and every second day with isotype control or NIMP-R14. Survival (upper panel) and weight loss (lower panel) was monitored daily (12 mice/group). **c-d**, Mice were treated as in 8b. Mice were euthanized on day 10 and lungs were collected. **c**, Individuals and means ± SEM of IFN-γ, IL-β and TNF-α levels in lung homogenates of isotype- or NIMP-R14-treated IAV-infected mice from two independent experiments are shown (7 mice/group). **d**, Individual values and means ± SEM of inflammatory monocytes, monocyte-derived DC, conventional DC and AM from two independent experiments are shown (6-8 mice/group). (**e-j**) IAV-infected mice were treated from 4 dpi and every second day with isotype control or anti-PD-L1 mAb. **e**, Survival was monitored daily (12 mice/group). (**f-h**), Mice were euthanized on day 10 and lungs were collected. **f**, Individuals and means ± SEM of IFN-γ, IL-β and TNF-α levels in lung homogenates of isotype- or anti-PD-L1-treated IAV-infected mice from two independent experiments are shown (6 mice/group). **g**, Individual values and means ± SEM of inflammatory monocytes, monocyte-derived DC, conventional DC, AM and classical neutrophils from two independent experiments are shown (8 mice/group). **h**, Unsupervised hierarchical clustering of the 30 genes imputed from transcriptional analysis of individual samples (n = 4/group). ns, not significant; *, p < 0.05; **, p < 0.01; ***, p<0.001.

As a complementary approach, IAV-infected mice were treated with a neutralizing anti-PD-L1 mAb (10F.9G2) starting at 4 dpi (**Fig. 8e**). In this setting, PD-L1 blockade also did not affect viral clearance (**Fig.S8c**) and led to higher local inflammation as judged by levels of inflammatory cytokines (**Fig. 8f**). PD-L1 neutralization led to an accumulation of MoDC and inflammatory monocytes (**Fig. 8g**). In addition, we observed a higher infiltration of PD-L1^-^ “classical” neutrophils upon this treatment (**Fig. 8g**). Again, this was paralleled with a significant decrease in AM absolute number (**Fig. 8g**). Moreover, we compared the transcriptional inflammatory signature of the lungs from control and anti-PD-L1-treated mice based on 30 selected genes encoding for proteins involved in IAV-related immune regulation. Unsupervised hierarchical clustering indicated the existence of three modules that could discriminate the two groups (**Fig. 8h**). In the lung of anti-PD-L1-treated mice, we noticed a reduced expression of genes encoding for factors associated with resolution of inflammation and tissue healing in IAV-induced pneumonia (module 2) as compared to controls such as *Areg*^39^, *Igf1*^40^*, Csf2*^41^, *Tgfb1*^42^*, Ptges2*^43^, *Il17rb*^44^ as well as genes encoding for angiogenic factors (*Vegfa*)^45^ (**Fig. 8h**). Conversely, module 1 comprised upregulated transcripts in PD-L1-treated mice for pro-inflammatory genes (*e.g. Ccl2*, *Ifng*, *Il13*) associated with deleterious IAV-mediated host response^46–48^ (**Fig. 8h**). Altogether, our findings suggest that PD-L1^+^ lung neutrophils control the deleterious inflammatory response associated with IAV infection.

## Discussion

Here, we demonstrated that lung viral infection led to the rapid emergence of regulatory PD-L1-expressing neutrophils within the BM in an IFN-γ-dependent manner and can be observed prior to their migration towards the inflamed/damaged tissue. In line with the literature^7^, we could detect PD-L1^+^ neutrophils in the blood of patients with severe VRI. Here, we also bring evidence for their high prevalence in airways of matched patients. VRI-associated PD-L1^+^ neutrophils were previously defined as “dysfunctional” with impaired antimicrobial activity^7^. Although they could be considered “dysfunctional” in regards to classical neutrophils, our data rather support that PD-L1^+^ neutrophils acquire a specific functional program associated with regulatory properties. While PD-L1^+^ neutrophils appear to exert a protective role in severe experimental VRI by limiting local inflammatory response, the situation in clinics is likely much more complex. Indeed, bacterial co-/super-infections often worsened the clinical picture during VRI^49^, and the emergence of regulatory neutrophils might also be at the expense of an optimal antimicrobial response^16^ and therefore might pave the way for these infections. Thus, this study identifies a pivotal, but sensitive, axis in the resistance/tolerance balance during severe viral pneumonia/ARDS. In clinics, neutrophils are still analyzed as a homogenous cell population, and further studies should be encouraged to better define the neutrophilic phenotype in relation to endotypes.

Emergency myelopoiesis arises to cope with an increased demand from the organism to replenish myeloid cells at the site of infection^17^. Our study supports the concept that emergency myelopoiesis may also constitute a demand-adapted response to deal with excessive inflammation and immunopathology^23,50^. Our observation is reminiscent with a population of monocytes that have been shown to be remotely educated by IFN-γ within the BM for regulatory functions upon *Toxoplasma gondii*-induced gut infection^50^. Therefore, sensing of IFN-γ by differentiating myeloid cells from multiple lineages could represent a broader regulatory mechanism during infections at mucosal surfaces. While IFN-γ is a prototypical inflammatory cytokine, it may also be part of a feedback circuit to limit immunopathology. The molecular and cellular actors that regulate this IAV-dependent IFN-γ response in the BM have not been explored. BM-resident innate (T) lymphocytes are interesting candidates as they were shown to produce IFN-γ upon innate signals^50,51^. It also remains unclear why only a fraction of BM neutrophils acquires a regulatory profile during experimental VRI. Although this may suggest the existence of a precommitted precursor, one can argue that the levels of IFN-γ are limiting in the BM during IAV infection. In line, the use of IFN-γ enabled to generate BMN with uniform regulatory phenotype.

Interestingly, the presence of a substantial proportion of PD-L1^+^ neutrophils in several non-lymphoid tissues (*e.g.* lungs, liver and gut (not shown)) -but not in spleen or lymph nodes (not shown) - under steady-state condition may suggest homeostatic functions in tissues exposed to numerous antigens, in which a high level of regulation and/or tolerance is required.

However, upon IAV infection, no increase in PD-L1^+^ neutrophils could be noted in non-lymphoid tissues, at the exception of the lungs, suggesting their preferential migration towards inflamed tissues. In line, BM PD-L1^+^ neutrophils from IAV-infected mice readily expressed the integrin CD49d^28^. Thus, BM PD-L1^+^ neutrophils appear to be equipped to migrate to inflamed tissue upon inflammation/infection, and therefore this may explain why the relatively low proportion of PD-L1^+^ neutrophils in the BM (∼10% of total neutrophils) can give rise to a large proportion of PD-L1^+^ neutrophils in the lungs of infected mice. We cannot rule out the contribution of other factors such as a high capacity to proliferate and/or the acquisition of the PD-L1^+^ phenotype once in the lungs. This latest hypothesis is nevertheless unlikely since lung PD-L1^-^ and PD-L1^+^ neutrophils presented highly divergent transcriptomes suggesting discrete sublineages. However, it is possible that additional signals received in the lungs can further tune PD-L1^+^ neutrophil functions. Indeed, our data suggest that the regulatory properties of PD-L1^+^ neutrophils varied during the course of IAV infection. For instance, the production of Arg-1 by lung PD-L1^+^ neutrophils is rather limited during the inflammatory phase and prevails during the resolution phase. This suggests that additional layers of regulation influence the functions of PD-L1^+^ neutrophils.

Moreover, their presence in naive mice in absence of strong IFN-γ signalling suggest the involvement of other factor(s) in the acquisition of the regulatory program. For instance, generation of BMN favours the PD-L1^+^ profile even in absence of IFN-γ, pointing towards other factor(s). Others factors such as G-CSF and TGF-β have been involved in the generation of PD-L1^+^ neutrophils with regulatory/suppressive functions upon thermal injury^16^ or cancer^52,53^. Interestingly, TGF-β-dependent expansion of PD-L1^+^ neutrophils occurs in the periphery and is transient (irrespective of cell death)^16^. Thus, it is likely that the biology of the regulatory/suppressive neutrophils may greatly varies according to the signals involved and tissue location. Further works into the licencing mechanisms leading to regulatory functions in neutrophils will undoubtedly offer new insights into how signals can be temporally and spatially coordinated and integrated to control the inflammatory balance. This is of particular interest during severe viral pneumonia/ARDS in which the therapeutic window has to be carefully considered for immunomodulatory approaches.

## Methods

### Clinical study design, patient population and approval

Over 18 yo patients with severe pneumonia and admitted in intensive care unit were prospectively included in this study, from January 2018 to May 2020. Only patients without diagnosed bacterial infections were enrolled. The clinical characteristics of the patients are documented in **Table 1**. The study was conducted in the University hospital of Tours (France). All patients or their next of kin gave their consent for enrolment in the study (ClinicalTrial.gov identifier: NCT03379207). This study was approved by the national ethic committee “Comité de Protection des Personnes Ile-de-France 8” under the agreement number 2017-A01841-52, in accordance with the French laws. Blood samples from healthy donors were obtained from the “Etablissement Français du Sang” (Agreement: CPDL-PLER-2019 188).

### Mice

Inbred and sex-matched 8-10 week-old C57BL/6j mice were purchased from Janvier (Le Genest-St-Isle, France), and maintained at the University of Tours under specific pathogen-free conditions. *Ifng*^-/-^ and control mice were maintained at the Pasteur Institute of Lille under specific pathogen-free conditions. *Ifngr1*^-/-^ mice were kindly provided by Dr. F. Laurent (INRAe, Nouzilly, France). All animal work was conformed to the French governmental and committee guidelines for the care and use of animals for scientific purposes; and was approved by the national ethic committee under approval number 201611151159949.

### Reagents and Abs

Monoclonal antibodies against mouse CD11b (M1/70), CD45 (30-f11), CD49d (R1-2), Sca-1 (D7), PD-L1/CD274 (10F.9G2), Arginase-1 (A1exF5), MHC-II (NIMR-4), Ly6G (1A8), CXCR4/CD184 (L276F12), F4/80 (BM8), IFN-γR1/CD119 (2E2), CD64 (X54-5/7.1), Ly6C (HK1.4) and Siglec F (E50-2440). Monoclonal antibodies against human CD16 (3G8), CD14 (M5E2), PD-L1/CD274 (29E.2A3) and CD10 (HI10a). All monoclonal antibodies and appropriated isotype controls were purchased from Biolegend (Amsterdam, The Netherlands), BD Pharmingen (Le Pont de Claix, France) and ThermoFisher Scientific/eBioscience (Paris, France). Dead cells were excluded with LIVE/DEAD® Fixable Aqua Dead Cell Stain kit (ThermoFisher Scientific, Illkirch, France). Mouse ELISA kits were from R&D systems (Lille, France) and eBioscience. Lineage Cell Depletion Kit was from Miltenyi-Biotec (Paris, France). Stem Cell Factor (SCF), IL-3 and Granulocyte-Colony Stimulating Factor (G-CSF) were purchased from Miltenyi-Biotec. Anti-mouse Ly6G/Ly6C (NIMP-R14) was kindly provided by Dr J.C. Sirard (Institut Pasteur de Lille, France). Purified anti-mouse CD3 (145-2C11) was from BD Pharmingen. Anti-mouse PD-L1 (clone 10F.9G2) and isotype controls were from Bio X Cell (Lebanon, NH, USA). Cell Trace™ Violet (CTV) cell proliferation kit was from ThermoFisher Scientific. Reparixin was from Sigma-Aldrich (Saint-Quentin-Fallavier, France). Recombinant mouse (rm)IFN-γ was purchased from ThermoFisher Scientific.

### IAV infection

The H3N2 IAV strain (A/Scotland/20/74) has been described elsewhere^54^. Mice were anesthetized and administered intranasally with 40 µl of PBS containing 150 plate-forming units. Weight and survival were monitored every second day following infection.

### Tissue harvest and preparation of cell suspensions

Lung cells were prepared as previously described^55^. Briefly, lungs were perfused with saline injected into the right ventricle. Lungs were harvested and minced using a gentleMACS dissociator (Miltenyi Biotec) in a medium containing 125 µg/ml of Liberase (Roche; Meylan, France) and 100 μg/ml of DNase type I (Roche). Lung homogenates were next resuspended in PBS and filtered onto a 100-µm cell strainer (Dutscher). Pellets were recovered in PBS 2% FCS and erythrocytes were removed using a red blood cell lysis buffer (Sigma-Aldrich) before being filtered onto a 40-µm cell strainer (Dutscher). Bronchoalveolar lavages (BAL) were performed after tracheal catheterisation by injecting four times 0,5 ml of cold PBS. BAL were then centrifuged at 400g for 5 min and pellets were recovered in PBS 2% FCS. Erythrocytes were removed using a red blood cell lysis buffer (Sigma-Aldrich) before being filtered onto a 40-µm cell strainer. For analysis of BM cells, femurs and tibias were collected and cut at their extremities, and cold RPMI was injected to harvest BM core biopsy, which was mechanically disrupted onto a 100-µm cell strainer. After centrifugation (400g for 5 min), erythrocytes were removed before being filtered onto a 40-µm cell strainer. For mRNA expression experiments, BM core biopsy were immediately frozen in liquid nitrogen. Blood was collected in microtubes filled with 30 µL of heparin from the retro-orbital sinus while mice were anesthetized. Erythrocytes were removed upon several washes in a red blood cell lysis buffer (Sigma-Aldrich).

For human neutrophil analyses, total blood was subjected to three repeated steps of erythrocyte lysis. Endotracheal aspirates (ETA) were collected from intubated patients and incubated with dithiothreitol (1 mM) in PBS (5 mL/g of ETA) for 30 min under continuous agitation. After centrifugation, pellet cells were filtered onto a 100-µM cell strainer. After removing red blood cells, cells were passed through a 40-µM cell strainer prior analysis.

### Oxidant production assay

Oxidant (RNS/ROS) production by neutrophils was assessed by flow cytometry using dihydro-rhodamine123 (DHR123). Lung homogenates were incubated with or without 100 nM PMA and with 1 µM DHR123 for 5 min at 37°C before analysis.

### *In vitro* neutrophil differentiation

Progenitors from the BM were enriched using the Lineage Cell Depletion Kit (Miltenyi). The negative fraction was then seeded into a 6-well plate at 1 x 10^5^ cell/ml in complete IMDM medium. Cells were differentiated for 7 days in presence of SCF (50 ng/mL), IL-3 (50 ng/ml) and G-CSF (50 ng/ml) as previously described^56^. In some cases, mouse recombinant IFN-γ (1 ng/ml) was added into the culture on day 0, 3 and 5.

### Proliferation assay

C57BL/6j mice were i.n. infected with IAV (150 pfu) and lungs were collected at 8 dpi. Neutrophils were then enriched using anti-Ly6G MicroBead kit (Miltenyi Biotec). In parallel, CTV-labelled spleen cells (5 × 10^5^) from naive mice were stimulated with plate-bound anti-CD3 mAb (4 µg/mL, clone 145-2C11) in 96-well culture plate in complete 10 % FCS RPMI1640 medium. Enriched neutrophils were then co-cultured with anti-CD3-stimulated spleen cells for 48 h in presence of Ig control (10 µg/mL, clone LTF-2) or anti-PD-L1 (10 µg/mL, clone 10F9G2) mAbs. Cells were then stained with anti-CD45, anti-CD3, anti-CD4, anti-CD8 antibodies and T cell proliferation rate was measured by monitoring loss of CTV fluorescence intensity.

### Flow cytometry

Cells were stained with appropriate dilutions of monoclonal antibodies. Then, cells were washed, and, in some cases, fixed and permeabilized using commercial kit from eBioscience according to the manufacturer’s information. Dead cells were excluded using the LIVE/DEAD Cell Staining kit. Cells were acquired on either a LSR Fortessa cytometer (BD Biosciences) or a MACS Quant (Miltenyi Biotec). Analyses were performed using the VenturiOne software (Applied Cytometry; Sheffield, UK).

### Cell sorting and *in vitro/ex vivo* assays

To purify neutrophil subsets, lung mononuclear cells were labelled with FITC-conjugated anti-Ly6G mAb, PerCp-Cy5.5-conjugated anti-CD11b mAb and APC-conjugated anti-PD-L1 mAb. After cell surface labelling, cells were sorted using a FACSMelody (BD Biosciences). This protocol yielded > 98 % cell purity. For *in vitro* stimulation assays, 5 x 10^4^ neutrophil subsets were cultured for 4 h in complete RPMI 5% FCS in absence of any additional stimulation. In some cases, neutrophils were pulsed for 5 min with PMA. Then, supernatants were harvested for further analyses and/or cells were subjected to cytometry analysis.

### Single-cell RNA-seq and data pre-processing

Single-cell suspension from 5 lungs of IAV-infected mice (day 11) were pooled and neutrophil subsets (LiveDead^-^ CD45^+^ SiglecF^-^ CD11b^+^ Ly6G^+^ PD-L1^+/-^) were sorted on a FACSMelody (BD Biosciences) (purity > 99%) into 1x PBS with 0.04% BSA + RNAse inhibitor. Sorted cells were counted under a microscope and 8,000-10,000 cells per subset were loaded onto a Chromium controller (10X Genomics). Reverse transcription and library preparation were performed according to the manufacturer’s protocol. Libraries were simultaneously sequenced on a NovaSeq 6000. Cell Ranger Single-Cell Software Suite v2.0.0 was used to perform sample demultiplexing, barcode processing and single-cell 30 gene counting using standards default parameters and *Mus musculus build mm10*. The argument “-force-cell” was used according to 10X genomics recommendations’ (https://support.10xgenomics.com/single-cell-gene-expression/software/pipelines/latest/tutorials/neutrophils).

### Single-cell RNA-seq analysis

#### Cell and gene filtering

Raw gene expression matrices generated per sample were passed to Seurat for preprocessing. Cells with less than 90 unique molecular identifiers (UMI) and more than 5 % of mitochondrial RNA were removed; then transcripts expressed in less than three cells were removed. Cells considered as doublet were also removed using DoubletFinder^57^. Thus, we could retain 1,165 cells from the PDL-1^-^ subset and 1,834 cells from the PD-L1^+^ subset. Concordance between datasets enabled to merge them for downstream analyses. Standard processing was used with an UMAP with k = 50 and min.dist = 0.3 using 20 principle components.

#### Trajectory analysis

Seurat dataset was converted into SingleCellExperiment to run trajectory analysis. All data from Monocle3 object were passed to Seurat.

#### Mapping gene along pseudotime

Pseudotime was calculated using Monocle3, pseudotime values were passed to Seurat object. Cells were ordered by pseudotime values. Spearman correlation was calculated for all genes with pseudotime. Top 20 highly positively and negatively correlated genes have been chosen to characterize the maturation of neutrophils.

#### Plots

All feature plots were created using scCustomize package. Density plots were made using Nebulosa package with “wkde” as kernel density estimation method^58^. ClusterMap was made using scanpy^59^. Gene ontology (GO) enrichment analysis were performed using ClusterProfiler2. The following GO datasets have been used: Regulation of gene expression (GO: 0010468), Regulation of cytoskeleton organization (GO: 0051493), Cell cycle process (GO: 0022402), Negative regulation of execution phase of apoptosis (GO: 1900118), Neutrophil-mediated antimicrobial response (GO: 0070944), Positive regulation of the inflammatory response (GO: 0050729), Negative regulation of the inflammatory response (GO: 0050728) IFN-γ-mediated signalling pathway (GO: 0060333). All gene sets can be found at https://www.gsea-msigdb.org/gsea/msigdb/mouse/genesets.jsp?collection=GO.

#### Data and Code availability

All data were submitted to the Gene Expression Omnibus accession number XXX. All R scripts used for single cell statistical analysis can be accessed on github (https://github.com/TEAM-4-CEPR/IFN-g-primes-bone-marrow-neutrophils-for-regulatory-functions-in-severe-viral-pneumonia).

### Analysis of gene transcripts by quantitative RT-PCR

RNAs from whole lungs or BM of naive or IAV-infected mice were extracted and purified using the NucleoSpin RNA Extraction Kit (Macherey-Nagel) according to the manufacturer’s instructions. cDNAs were synthesized using High Capacity RNA-to-cDNA kit (ThermoFischer Scientific) and quantitative RT-PCR were carried out using specific primers (**Table S2**) and SYBR Green PCR Master Mix (Qiagen). PCR amplification of *Gapdh* was performed to control for sample loading and normalization between samples. ΔCt values were obtained by deducting Ct values of *Gapdh* mRNA from the Ct values obtained for the investigated genes. Data are represented as relative expression of mRNA levels.

### Neutrophil depletion and PD-L1 blockade

Neutrophils were depleted using NIMP-R14 (100 μg per mouse i.p. in 200 µl PBS) every second day at the indicated starting point. 10F.9G2 was injected (200 μg per mouse i.p. in 200 µl PBS) every second day from 4 dpi. A rat IgG2b (LTF2) and a mouse IgG2b (MPC-11) isotype control antibodies were used as controls.

### Statistical analysis

All statistical analysis was performed using GraphPad Prism software. The statistical significance was evaluated using non-parametric Mann-Whitney U tests or Kruskal-Wallis to compare the means of biological replicates in each experimental group. Survival rates were analyzed using a log-rank test. Results with a P value of less than 0.05 were considered significant. ns: not significant; *, p < 0.05; **, p <0.01; ***, p < 0.001.

## Abbreviations

ARDS: acute respiratory distress syndrome;
dpi: days post-infection;
IAV: influenza A virus;
MHC: major histocompatibility complex;
NET: neutrophil extracellular trap;
PD-L1: Programmed death-ligand 1;
VRI: viral respiratory infection;
WT: wild-type.

## Acknowledgements

We thank the University of Tours (PST-A) and the Institut Pasteur de Lille animal facilities for their excellent mouse husbandry. The Cytometry and Single-cell Immunobiology core facility (University of Tours, PST-ASB) is also acknowledged for technical assistance. We are grateful to all healthcare co-workers involved especially Aurélie Aubrey, Delphine Chartier and Véronique Siméon for their excellent management of patient samples and clinical data. Stephan Ehrmann, Pierre-François Dequin, Annick Legras, Denis Garot, Emmanuelle Mercier, Charlotte Salmon-Gandonnière, Laetitia Bodet-Contentin, Marlène Morisseau, Stephan Mankikian and Walid Darwiche are acknowledged for patient inclusions. We thank all the patients and their families for their trust and confidence in our work.

## Author contributions

Conceptualization: YJ, TM, CF, TB, CP

Methodology: FC, LG, EB, GI, TM, CF, TB, CP

Investigation: FC, YJ, LG, EB, RL, DS, CB, QL, CdAH, VS, TB, CP

Analysis: FC, YJ, LG, EB, GI, RL, QL, CdAH, TB, CP

Resources: AG, NW, FT, MS, BB, TM, CF, TB, CP

Funding acquisition: CP

Supervision: CP

Writing – original draft: CP

Writing – review & editing: YJ, LG, EB, GI, RL, AG, NW, FT, MS, BB, TM, CF, TB, CP

