## Supplementary information for "IFN-γ primes bone marrow neutrophils to acquire regulatory functions in severe viral respiratory infections"

**Supplementary Figures**

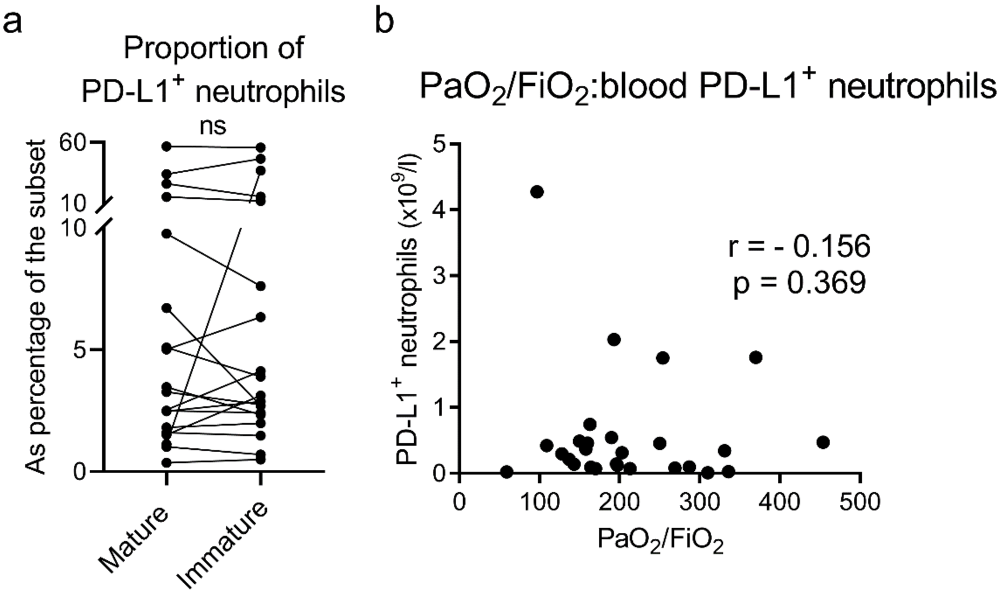


**Figure S1: PD-L1⁺ neutrophils in blood of patients with severe VRI. a,** Proportion of blood PD-L1+ neutrophils in mature vs immature subsets. **b**, Spearman's rank correlation of blood PD-L1⁺ neutrophils and hypoxemia levels on admission of intubated patients with viral pneumonia (n =27). Ns, not significant.

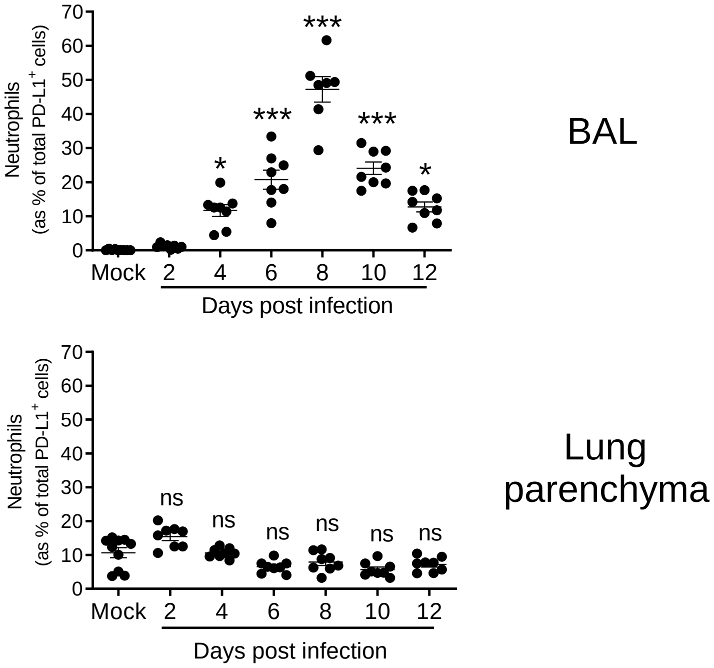


**Figure S2: Frequency of neutrophils within total PD-L1⁺ leukocytes.** Relative proportion of neutrophils within total PD-L1⁺ leukocytes in airways and lung parenchyma was evaluated by flow cytometry. Individual and means ± SEM pooled from three independent experiments are shown in the right panel (8-10 mice/group). ns, not significant; *, $p < 0.05$; ***, $p < 0.001$.

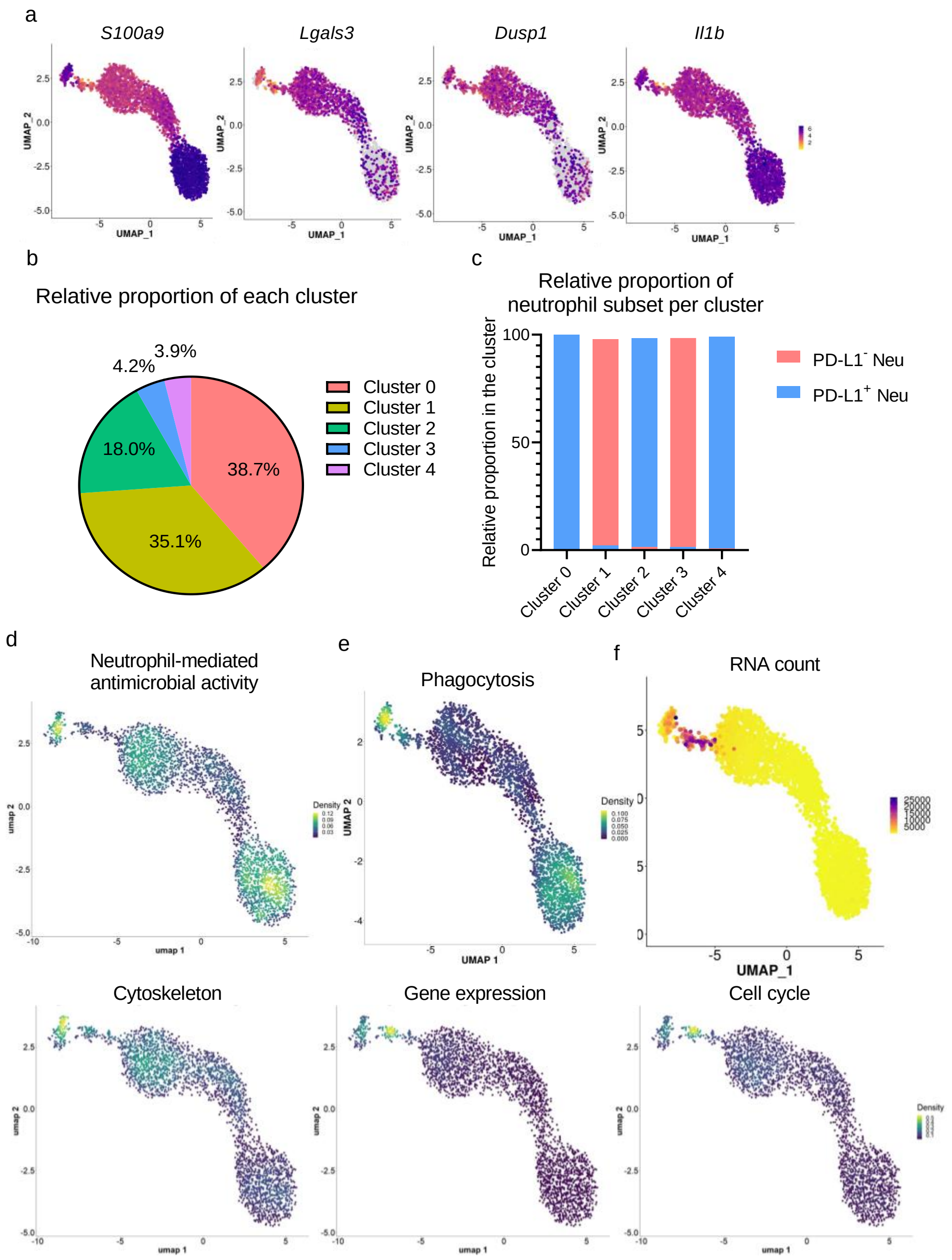


**Figure S3: scRNA-seq of lung neutrophilic subsets based on PD-L1 expression in IAV-infected C57Bl/6j mice. a**, Expression map of selected neutrophil gene markers. **b**, Relative proportion of each cluster in merged data set. **c**, Relative proportion of PD-L1$^{-}$ and PD-L1$^{+}$ subsets in each defined cluster. **d**, "Neutrophil-mediated antimicrobial activity" signature along merged data. **e**, "Phagocytosis" signature along merged data. **f**, Number of reads per cell along merged data **g**, Expression of signature from cytoskeleton, gene expression (nFeatures) and cell cycle.

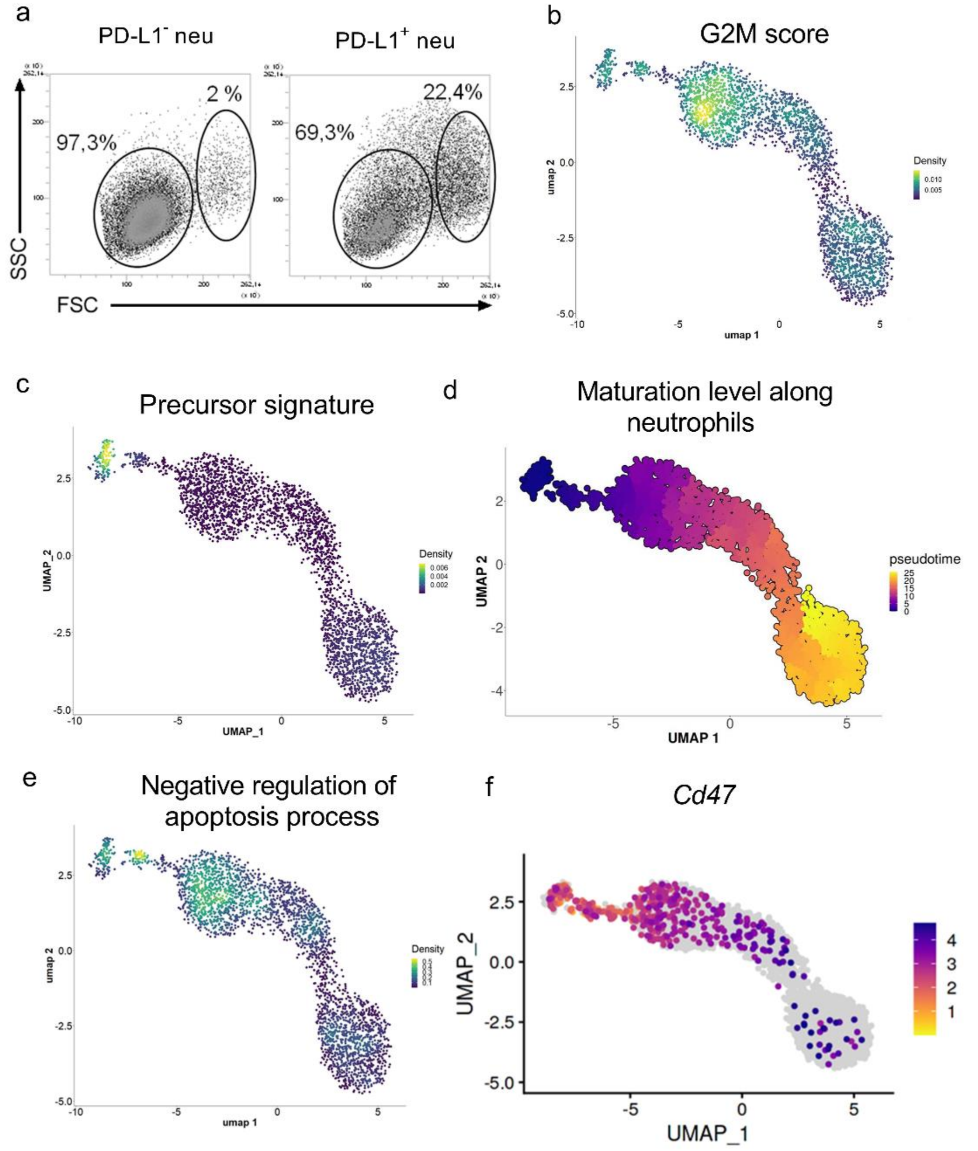


**Figure S4: Proliferative, immature and anti-apoptotic transcriptional profile of lung PD-L1$^{+}$ neutrophils in IAV-infected C57Bl/6j mice**. **a,** Representative dot plots of FSC/SSC of lung neutrophil subsets from IAV-infected mice. **b,** Density plot of G2M cell cycle score were extracted from Seurat package. **c**, Density plot of the “precursor” signature in neutrophil transcriptomes. **d**, Maturation level mapped with pseudotime along merged data with precursor signature used as root. **e**, “Negative regulation of apoptosis process” signature in neutrophil transcriptomes. **f**, Expression map of *Cd47* in neutrophil transcriptomes

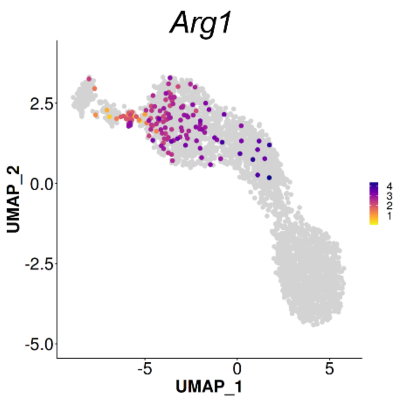


**Figure S5: Lung PD-L1$^{+}$ neutrophils from IAV-infected C57Bl/6j mice expressed *Arg1*.** Expression map of *Arg1* in neutrophil transcriptomes from **Fig. 3**.

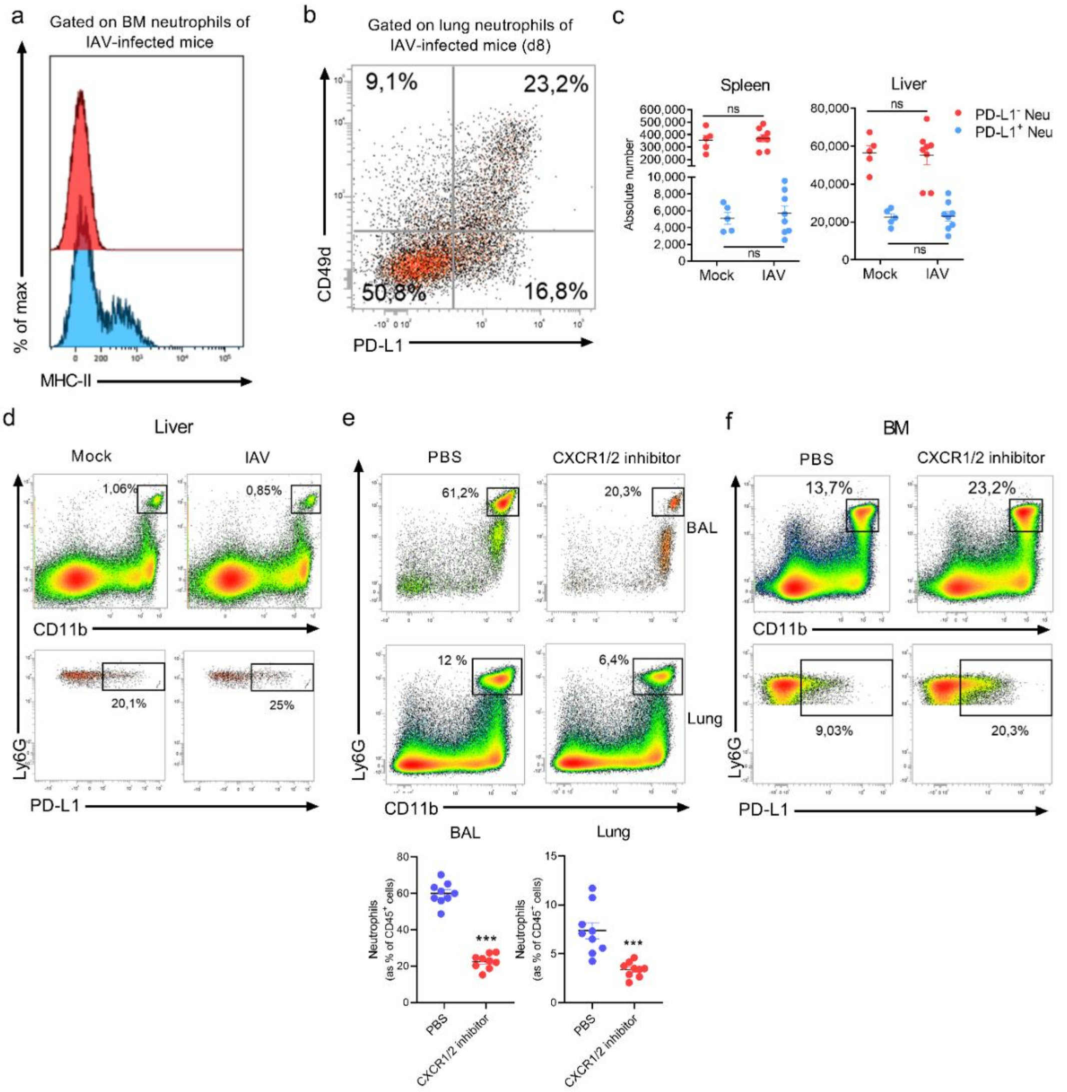


**Figure S6: Origin and migration of IAV-induced PD-L1⁺ neutrophils in C57Bl/6j mice.** **a**, Flow cytometry expression of MHC-II on BM neutrophil subsets. Representative dot plots of 4 independent experiments of PD-L1⁻ and PD-L1⁺ BM neutrophils from mock or IAV-infected mice are shown. **b**, Flow cytometry expression of CD49d on lung neutrophils from IAV-infected mice according to PD-L1 expression. Representative dot plots of 3 independent experiments are shown. **c**, Absolute number of neutrophil subsets in spleen and liver of mock and IAV-infected mice. Individual values and means ± SEM from two independent experiments are shown (n = 5-8/group). **d**, Proportion of PD-L1⁺ neutrophils in the liver of mock and IAV-infected mice. Representative dot plots of 3 independent experiments are shown. **e-f**, Proportion of neutrophils subsets in IAV-infected mice treated or not with reparixin (CXCR1/2 inhibitor). **e**, Neutrophil proportions in BAL and lung parenchyma of IAV-infected mice treated or not with reparixin. Representative dot plots from two independent experiments are shown in the left panel. Individual values and means ± SEM from two independent experiments are shown (n = 9/group) in the right panel. **f**, Neutrophil proportions in BM of IAV-infected mice treated or not with reparixin. Representative dot plots from two independent experiments are shown. ns, not significant; ***, $p < 0.001$.

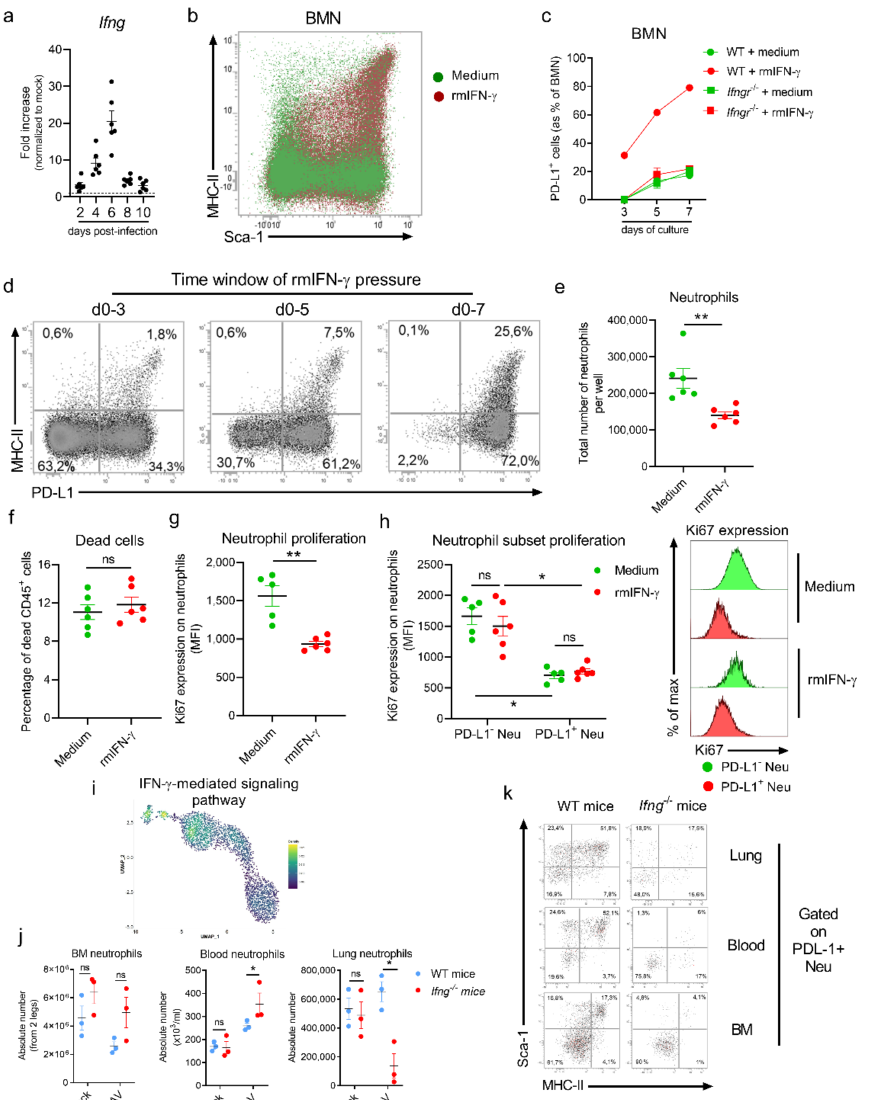


Figure S7

**Figure S7: Influence of IFN-γ on emergence of BM PD-L1⁺ neutrophils. a**, *Ifng* mRNA expression in BM was determined by RT-qPCR. Data are normalized to expression of *Gapdh* and are expressed as fold increase over average gene expression in mock-treated mice. Individual values and means ± SEM from two independent experiments are shown (n = 6/group). **b**, Flow cytometry expression of Sca-1 and MHC-II on BMN differentiated in presence or not of rmIFN-γ. Representative overlay dot plots are shown. **c**, Kinetic representation of the proportion of PD-L1$^+$ BMN differentiated from WT or *Ifngr*$^{-/-}$ BM stem cells in presence or not of rm-IFN-γ. Means ± SEM from two independent experiments are shown. **d**, Representative dot plots showing PD-L1 and MHC-II expression on differentiated BMN according to the length of rm-IFN-γ are shown. **e**, Effect of rm-IFN-γ on the absolute number of BMN obtained at the end of the differentiation protocol. Individual values and means ± SEM from two independent experiments are shown. **f**, Relative proportion of dead cells at the end of the protocol defined using the Live/Dead stain kit. Individual values and means ± SEM from two independent experiments are shown. **g**, Flow cytometry expression of the proliferation marker Ki67 in differentiated BMN in presence or not rm-IFN-γ. Individual values and means ± SEM from two independent experiments are shown. **h**, Differential proliferation rate in PD-L1$^-$ vs PD-L1$^+$ BMN measured by Ki67 expression. Individual values and means ± SEM from two independent experiments are shown. Representative overlay histograms are shown in the right panel. **i**, IFN-γ-mediated signalling pathway signature in neutrophil transcriptomes. **j**, Absolute numbers of neutrophils in BM, blood and lung of IAV-infected WT or *Ifng*$^{-/-}$ mice. **k**, Representative dot plots of one experiment out of two of Sca-1 and

MHC-II expression in lung, blood and BM PD-L1+ neutrophils from IAV-infected WT or *Ifng*-/- mice. ns, not significant; *, p < 0.05; **, p < 0.01.

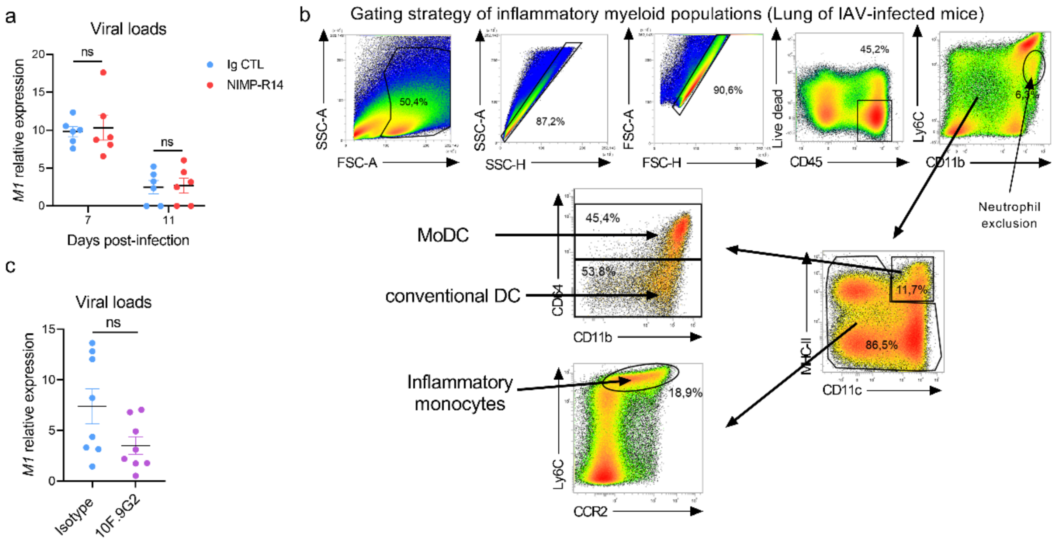


**Figure S8: Viral loads and gating strategy of myeloid cell subsets in lung of IAV-infected mice. a**, Analysis of the viral load in the lung of IAV-infected WT mice treated with Ig control or NIMP-R14. IAV *M1* mRNA relative expression in the whole lung were measured by quantitative RT-PCR on the indicated day. Individual values and means ± SEM from two independent experiments are shown. **b**, gating strategy of the lung inflammatory myeloid subsets. Representative dot plots of one experiment out of 6 are shown. **c**, Analysis of the viral load in the lung of IAV-infected WT mice treated with Ig control or 10F.9G2. IAV *M1* mRNA relative expression in the whole lung were measured by quantitative RT-PCR. Individual values and means ± SEM from two independent experiments are shown. ns, not significant.
