## Supplementary material for "IFN-γ primes bone marrow neutrophils to acquire regulatory functions in severe viral respiratory infections": Table 1

| | **Overall** (n=35) | **COVID-19** (n=27) | **Influenza** (n=8) |
|---|---|---|---|
| Age (year), median (IQR) | 64 (60; 71) | 64 (59; 70) | 66 (61; 74) |
| Male / female, n/n (%) | 21/14 (60%) | 17/10 (63%) | 4/4 (50%) |
| BMI (kg/m$^2$), median (IQR) | 32 (29; 35) | 31 (28;25) | 33 (31; 36) |
| Type 2 diabetes, n (%) | 10 (30%) | 8 (30%) | 2 (25%) |
| Hypertension | 17 (49%) | 13 (48%) | 4 (50%) |
| Chronic respiratory disease, n (%) | 1 (3%) | 0 | 1 (13%) |
| Chronic kidney disease, n (%) | 2 (6%) | 1 (4%) | 1 (13%) |
| Chronic cardiovascular disease, n (%) | 0 | 0 | 0 |
| SAPS2, median (IQR) | 34 (25; 42) | 32 (23; 39) | 46 (37; 55) |
| SOFA at inclusion, median (IQR) | 4 (3; 6.5) | 4 (2; 6) | 8.5 (4; 10) |
| Invasive mechanical ventilation at inclusion, n (%) | 26 (74%) | 19 (70%) | 7 (88%) |

**Table 1. Patients' baseline characteristics at inclusion**. IQR : interval quartile range ; BMI : Body Mass Index, SAPS2 : Simplified Acute Physiology Score 2 ; SOFA : Sequential Organ Failure Assessment.
